# Transcriptional impacts of substance use disorder and HIV on human ventral midbrain neurons and microglia

**DOI:** 10.1101/2025.02.05.636667

**Authors:** Alyssa M. Wilson, Michelle M. Jacobs, Tova Y. Lambert, Aditi Valada, Gregory Meloni, Evan Gilmore, Jacinta Murray, Susan Morgello, Schahram Akbarian

**Author notes:** Albert Einstein College of Medicine, New York, New York 10461, USA. These authors contributed equally to this work (Susan Morgello, Schahram Akbarian).

## Abstract

For people with HIV (PWH), substance use disorders (SUDs) are a prominent neurological risk factor, and the impacts of both on dopaminergic pathways are a potential point of deleterious convergence. Here, we profile, at single nucleus resolution, the substantia nigra (SN) transcriptomes of 90 postmortem donors in the context of chronic HIV and opioid/cocaine SUD, including 67 prospectively characterized PWH. We report altered microglial expression for hundreds of pro- and anti-inflammatory regulators attributable to HIV, and separately, to SUD. Stepwise, progressive microglial dysregulation, coupled to altered SN dopaminergic and GABAergic signaling, was associated with SUD/HIV dual diagnosis and further with lack of viral suppression in blood. In virologically suppressed donors, SUD comorbidity was associated with microglial transcriptional changes permissive for HIV infection. We report HIV-related downregulation of monoamine reuptake transporters specifically in dopaminergic neurons regardless of SUD status or viral load, and additional transcriptional signatures consistent with selective vulnerability of SN dopamine neurons.

## Introduction

The HIV pandemic continues to pose a global health burden, despite considerable advances in treatment and prevention^1,2^. Although a large fraction of people with HIV (PWH) are receiving life-saving antiretroviral therapy (ART), treatment does not eradicate the virus, and HIV persists in multiple tissue reservoirs including the brain. PWH may develop neurologic disorders that are predicated in part on immune dysfunction, and that are presumed to result from the action of HIV on nervous system cells^2–4^. An important co-morbidity relevant to HIV neurologic disease is substance use disorder (SUD), and in particular injection drug use^5^, with its negative impact on health by a multitude of factors including risky behaviors, emergence of medical and psychiatric sequelae, and lack of compliance with therapy^6–15^. Notably, HIV and SUD show a striking neuroanatomical convergence on dopamine-rich structures in brain. In ventral midbrain^16^ this includes substantia nigra (SN)^17,18^, a region important for mediating habitual behaviors and salience of cues associated with drug use^18^, withdrawal-related anhedonia and dysphoria, and extra-pyramidal motor dysfunction^19,20^. There is also evidence, mostly from preclinical work, that drug-of-abuse-induced dynamic adaptations in addiction-relevant dopaminergic circuitry could worsen HIV-associated cognitive and behavioral deficits^21^, with multiple dopaminergic signaling pathways converging on susceptible myeloid cells to promote viral entry and replication^22,23^.

Despite its importance, very little is known about cellular and molecular alterations in the ventral midbrain of PWH with or without SUD comorbidity, with the majority of studies limited to quantification of neurotransmitter (e.g., dopamine) in bulk tissue or neuroimaging and analysis of cerebrospinal fluid (CSF) in the context of advanced disease (AIDS)^24–26^. Herein, we profile for the first time, the ventral midbrain transcriptomes from n=90 total PWH and people without HIV (“HIV-”), at single (cell) nucleus resolution, and deliver first insights into the cell-type-specific neurogenomic responses to chronic infection and SUD. We specifically examine responses to opioid and/or cocaine SUDs, given the high prevalence of these substances and their combination among PWH^27–29^. Our analyses, focused on SN dopaminergic (DA) and GABAergic (GABA) neurons and their surrounding microglia, strongly suggest that SUD for opioids and cocaine creates, even in donors with antiretroviral therapy (ART)-mediated viral suppression, a molecular environment permissive for HIV viral replication and risk for cytotoxic damage to dopaminergic neurons. In addition, we report a stepwise progression of transcriptomic dysregulation for opioid/cocaine SUD and HIV comorbidity that culminates in widespread neuronal pathology and pronounced inflammatory signatures in microglia from individuals without viral suppression (viremia) in the context of SUD.

## Results

### SN cell type composition in the context of HIV and SUD

We dissected ventral midbrain for unilateral collection of the SN including *pars compacta* and *pars reticulata* from n=90 donors collected by the Manhattan HIV Brain Bank (MHBB) (39 female/ 51 male; mean+/-standard deviation age 54.0+/-11.4 years; 33% Hispanic, 38% non-Hispanic Black, 22% non-Hispanic White; Table 1), including 67 participants followed longitudinally in the MHBB with deep clinical phenotyping during years of study participation (median 6, maximum 21 years). We assigned each donor to one of three HIV statuses: negative (“HIV-”, n=28 donors), HIV positive with “undetectable” final plasma viral load (VL) defined as <50 RNA copies/mL (“HIV+u”, n=30 donors), or HIV positive with viremia, evidenced by a detectable final plasma VL (“HIV+d”, n=32 donors [viral RNA copies/mL range 65 to >750,000, median 6,721]). We further grouped donors by substance use into “SUD-” or “SUD+” for any combination of opioid/cocaine SUD (n=60 SUD+, including n=42 with a history of misusing opioids, of which n=28 had a history of misusing both opioids and cocaine). This grouping produced HIV-/SUD-defined subgroups that were overall closely matched with regard to demographic factors: SUD-HIV-(n= 13); SUD+HIV-(n=15); SUD-HIV+u (n=10); SUD+HIV+u (n=20); SUD-HIV+d (n=7); and SUD+HIV+d (n=25) (Fig. 1A, Tables S1-S3). Notably, the wider MHBB cohort from which these donors were selected had a high prevalence of polysubstance SUD, with approximately three quarters of SUD-positive individuals endorsing symptoms for two or more drug classes.

**Table 1.** Demographic characteristics of donor groups.

|  | **Entire Sample** | **SUD-HIV-**<br>**(n=13)** | **SUD+HIV-**<br>**(n=15)** | **SUD-HIV+u**<br>**(n=10)** | **SUD+HIV+u**<br>**(n=20)** | **SUD-HIV+d**<br>**(n=7)** | **SUD+HIV+d**<br>**(n=25)** |
| --- | --- | --- | --- | --- | --- | --- | --- |
| **Mean (standard deviation) age [years]** | 54.0 (11.4) | 47.5 (2.9) | 54.3 (2.7) | 60.1 (3.3) | 60.8 (2.3) | 54.0 (4.0) | 49.4 (2.1) |
| **Number (percent) male sex** | 51 (57%) | 7 (54%) | 12 (80%) | 5 (50%) | 10 (50%) | 4 (57%) | 13 (52%) |
| **Race/ethnicity by number (percent)** | 34 (38%)<br>Non-Hisp<br>Black | 2 (15%)<br>Non-Hisp<br>Black | 4 (27%)<br>Non-Hisp<br>Black | 2 (20%)<br>Non-Hisp<br>Black | 9 (45%)<br>Non-Hisp<br>Black | 3 (43%)<br>Non-Hisp<br>Black | 14 (56%)<br>Non-Hisp<br>Black |
|  | 30 (33%)<br>Hisp. | 6(46%)<br>Hisp | 5(33%)<br>Hisp | 2(20%)<br>Hisp | 8(40%)<br>Hisp | 1(14%)<br>Hisp | 8(32%)<br>Hisp |
|  | 20 (22%)<br>Non-Hisp<br>White | 3 (23%)<br>Non-Hisp<br>White | 6 (40%)<br>Non-Hisp<br>White | 6 (60%)<br>Non-Hisp<br>White | 1 (5%)<br>Non-Hisp<br>White | 1 (14%)<br>Non-Hisp<br>White | 3 (12%)<br>Non-Hisp<br>White |
|  | 6 (7%)<br>Other | 2 (15%)<br>Other | 0 (0%)<br>Other | 0 (0%)<br>Other | 2 (10%)<br>Other | 2 (29%)<br>Other | 0 (0%)<br>Other |
| **Median (IQR) CD4 T cell count** | 165 (71, 425) | N/A | N/A | 378 (208, 648) | 269 (135, 421) | 242 (185, 400) | 69 (15, 102) |
| **Median (IQR) nadir CD4 T cell count** | 54 (12, 139) | N/A | N/A | 113 (80, 270) | 60 (13, 172) | 93 (17, 185) | 22 (6, 58) |
| **Opioid/cocaine SUD diagnoses by number** | Opioid and cocaine<br>28 | N/A | Opioid and cocaine<br>10 | N/A | Opioid and cocaine<br>10 | N/A | Opioid and cocaine<br>8 |
|  | Opioid only<br>14 |  | Opioid only<br>4 |  | Opioid only<br>4 |  | Opioid only<br>6 |
|  | Cocaine only<br>18 |  | Cocaine only<br>1 |  | Cocaine only<br>6 |  | Cocaine only<br>11 |
| **Number (percent) with exposure to other substances** | 67 (74%) | 3 (23%) | 10(67%) | 8 (80%) | 19 (95%) | 6 (86%) | 21 (84%) |

Statistical comparisons of donor characteristics across donor groups are provided in Table S2.
Numbers of nuclei by donor group are provided in Table S3.

**Fig. 1.**
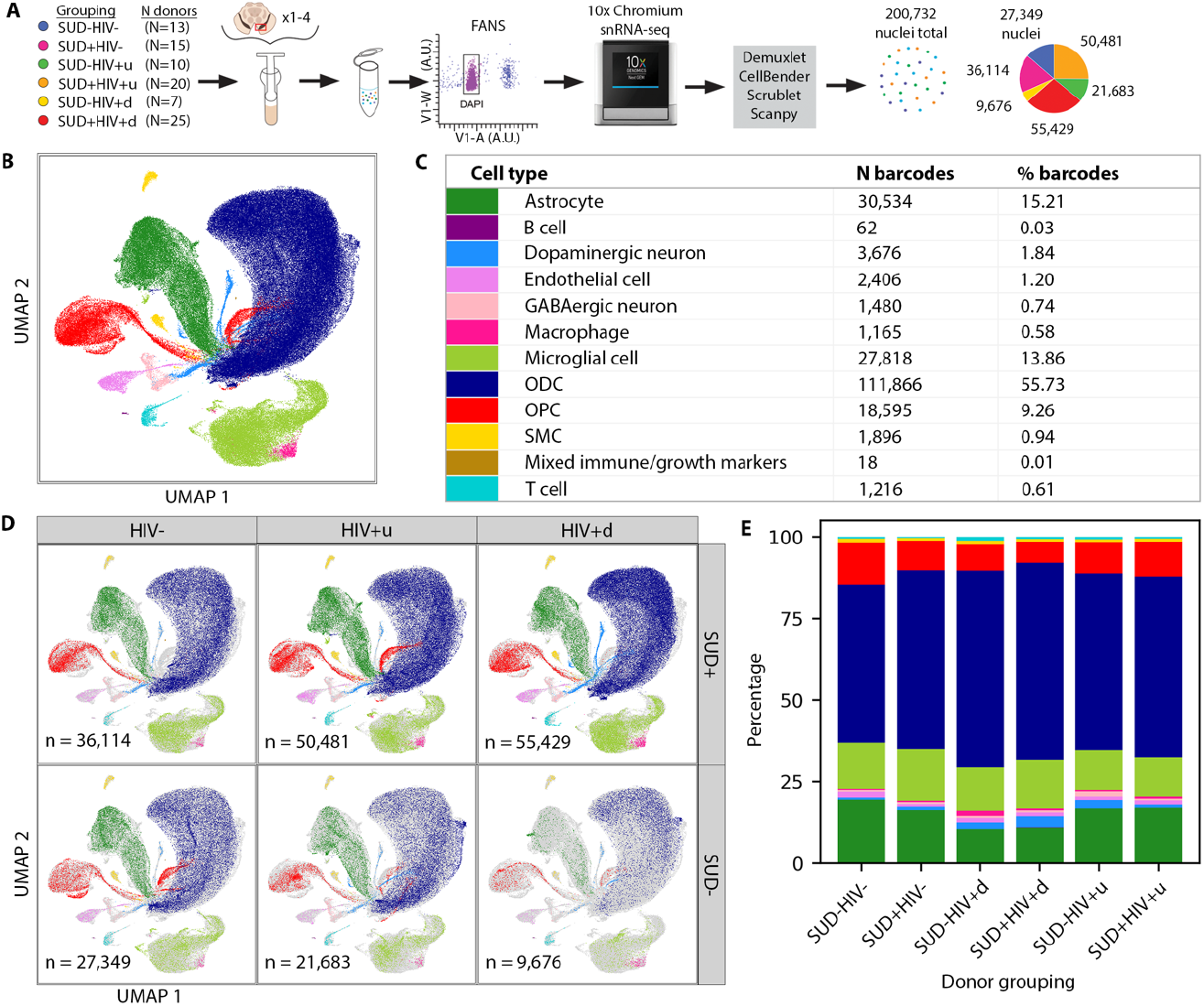
snRNA-seq data generation and cell typing results. **(A)** Schematic of study design and snRNA-seq data acquisition procedure (showing at left the number of donors per study group and at right, with the same color-coding, each group’s final number of QC-passing nuclei). “V1-W” and “V1-A”: width and area, respectively, of violet-channel fluorescence signal used for detecting intact nuclei with fluorescence-activated nuclear sorting (FANS); “DAPI”: 4′,6-diamidino-2-phenylindole. **(B)** 2D UMAP showing the cell types we identified, with cell type colors as shown in **(C). (C)** Numbers and fractions of nuclei of each cell type. **(D)** UMAPs showing nuclei by cell type for each donor group (colors as in **(C)**). **(E)** Cell type proportions for each donor group (see also Table S5), with colors as shown in **(C)**: red, oligodendrocyte precursor cells (OPC); dark blue, oligodendrocytes (ODC); light green, microglia; dark green, astrocytes; and less-frequent cell types including in light blue, DA neurons and rose, GABA neurons.

We processed nuclei in pools of up to 4 donors, with most pools containing mixed HIV/SUD donor phenotypes, using the 10x Chromium 3’ system followed by Illumina sequencing. We then matched each single, sequenced nucleus to a specific donor using SNP-array genotyping and demuxlet^30^, and applied a series of quality control (QC) filters to each nucleus (Figs. S1A-C), including an upper threshold on the proportion of mitochondrial reads (set at 0.0128; Fig. S1A). We note that 100% of glial and 98% of neuronal nuclei that passed QC had a minimum of 200 genes expressed (Figs. S1B, C), thereby exceeding standards from published SN reference sets^31,32^ (compare Figs. S2A and S2B). After preprocessing, the final dataset amounted to a total of 200,732 QC-passing nuclei, with 81/90 donors contributing >1,000 nuclei (Fig. S1D; Table S4; Methods).

We first asked whether SN cell type composition is affected in the context of HIV and SUD. To maximize sensitivity for potential shifts in cellular subpopulations, we used as input 20,000/36,601 gene transcripts (Fig. S1E) as opposed to more typically suggested sizes of ∼2,000^33^, and we performed a likelihood-based, agnostic approach for cell typing (Methods). Our approach produced 12 major cell types (Figs. 1B, C, Fig. S3) that by numbers and proportions closely matched those reported for human SN single-cell-based transcriptomic reference sets^34,35^, with the largest share (65%) composed of oligodendrocytes and their precursors, and 14-15% made up of microglia and astrocytes. In contrast, SN neurons, endothelium, lymphocytes and macrophages each contributed a much smaller proportion of nuclei. There were no significant differences for 176/180 statistical comparisons of cell type proportions across HIV/SUD donor groups, indicating that the overall representation of major cell types in SN is preserved independent of HIV or SUD status (Figs. 1D, E; Table S5).

### Cell-specific transcriptomic responses to HIV infection show progression in the context of SUD, as well as SUD-independent downregulation of monoamine transporters in DA neurons

To examine cell-level impacts on SN function due to SUD, HIV, or both, we performed differential expression analysis (DEA; Methods) by stratifying the entire cohort of n=90 SN samples first by HIV groupings to interrogate effects of SUD, thereby comparing SUD+ vs. SUD-for HIV-donors, SUD+ vs. SUD-for HIV+u donors, and SUD+ vs. SUD-for HIV+d donors (Fig. 2). Likewise, to examine the impacts of HIV status, we stratified by SUD grouping, comparing HIV+u vs. HIV-for SUD+ donors, then for SUD-donors, and repeating similar analyses for HIV+d vs. HIV- and HIV+d vs. HIV+u, first for SUD+ and then for SUD-donors (Figs. 3-5).

**Fig. 2.**
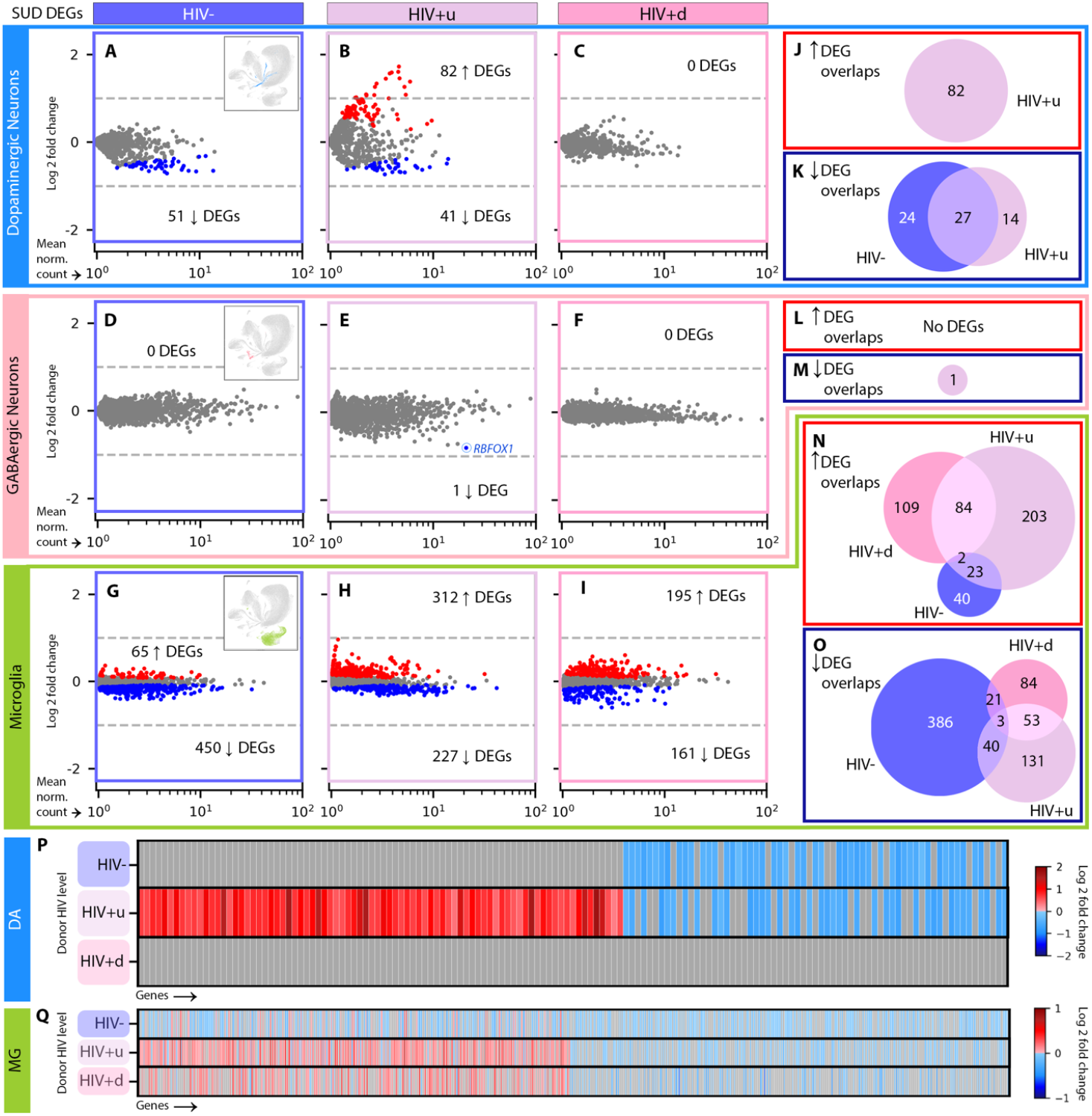
SUD-attributed DEGs by cell type. **(A)-(C)** SUD DEGs in DA neurons of **(A)** HIV-(dark purple), **(B)** HIV+u (light purple), and **(C)** HIV+d (pink) donors. Red and blue dots: significantly up- and down-regulated DEGs, respectively (p < 0.05). Numbers of up- and down-DEGs are shown. **(D)-(F):** SUD DEGs in GABA neurons for **(D)** HIV-, **(E)** HIV+u, and **(F)** HIV+d donors. In **(E)**, the DEG *RBFOX1* is circled. **(G)-(I)** SUD DEGs in microglia for **(G)** HIV-, **(H)** HIV+u, and **(I)** HIV+d donors. **(J)-(O)** Venn diagrams showing the extent of overlap among SUD DEGs across HIV levels for **(J)** DA neuron up- and **(K)** down-DEGs; **(L)** GABA neuron up- and **(M)** down-DEGs; and **(N)** microglia up- and **(O)** down-DEGs. **(P)-(Q)** Gene-by-gene comparisons of differential transcription across HIV levels for **(P)** DA neurons and **(Q)** microglia. Columns in **(P), (Q)** contain all genes arising as SUD DEGs at any HIV level; rows correspond to HIV levels. Genes per HIV level (row) are colored by log2 fold change (l2fc) if they are DEGs (see color bar) or gray if not. DA neuron, GABA neuron, and microglial nuclei are shown in the UMAP insets in **(A), (D)**, and **(G)**, respectively.

**Fig. 3.**
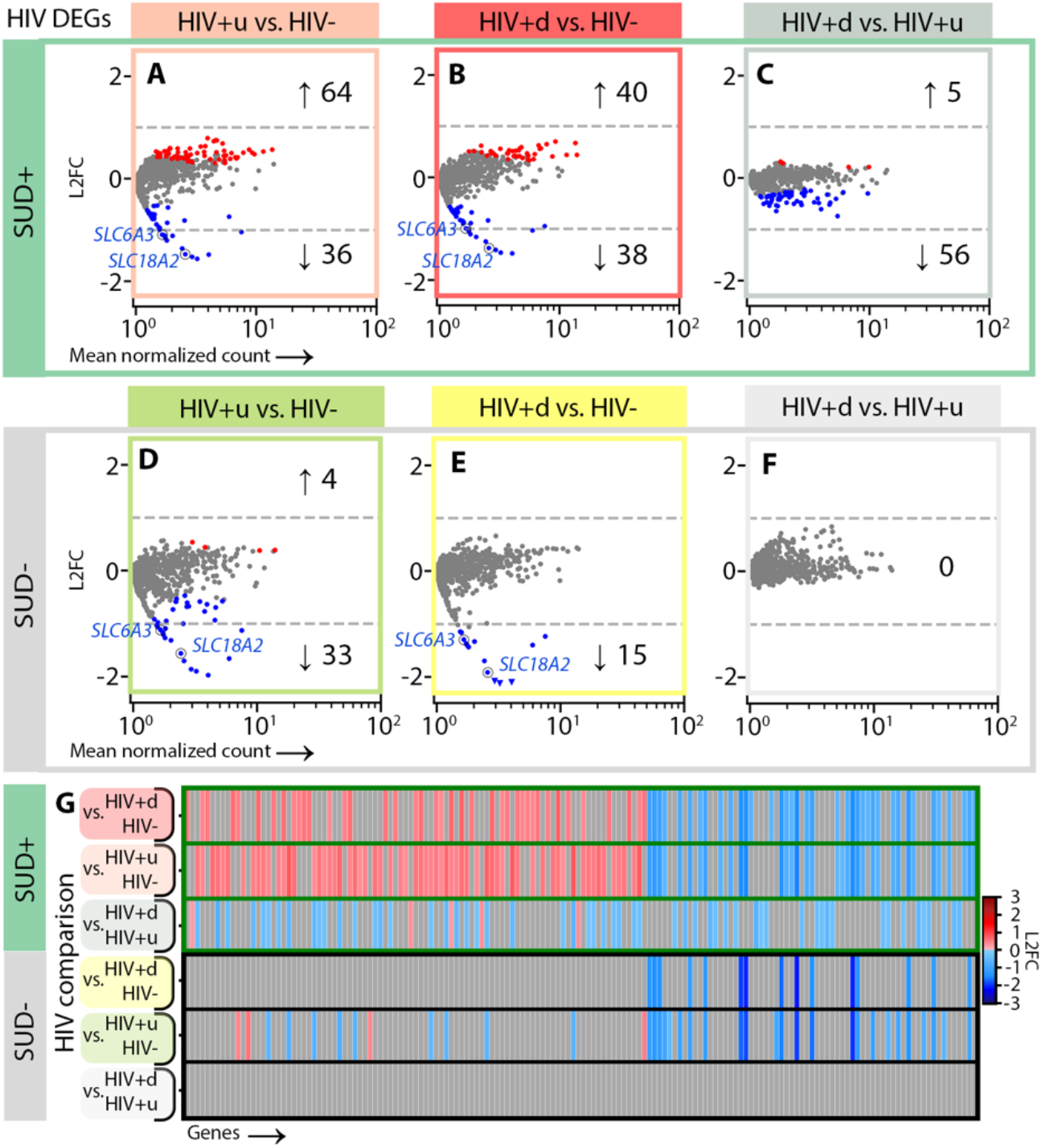
HIV-attributed DEGs for DA neurons. **(A)-(C)** HIV DEGs occurring in SUD+ individuals (p < 0.05) for DA neurons, for DEAs of **(A)** SUD+HIV+u vs. SUD+HIV- (peach), **(B)** SUD+HIV+d vs. SUD+HIV- (red), and **(C)** SUD+HIV+d vs. SUD+HIV+u (gray). **(D)-(F)** HIV DEGs occurring in SUD-individuals for DEAs of **(D)** SUD-HIV+u vs. SUD-HIV- (green), **(E)** SUD-HIV+d vs. SUD-HIV- (yellow), and **(F)** SUD-HIV+d vs. SUD-HIV+u (gray). **(G)** Gene-by-gene comparisons of differential transcription across HIV comparisons for (top 3 rows) SUD+ and (bottom 3 rows) SUD-, similar to **Figs. 2P, Q.** In **(A), (B), (D)**, and **(E)**, monoamine transporter DEGs *SLC6A3* and *SLC18A2* are circled.

We focused our DEAs on SN DA neurons and GABA neurons, given their key role in addiction circuitry and potential sensitivity to HIV infection^25,36–39^, and also on microglia, as mediators of neuroinflammation and host cells for HIV genomic integration^39–41^. We reduced the rate of false-positive differentially expressed genes (DEGs) using multiple steps. First, we applied cross-fold validation (*k*=3 folds), creating for each comparison and cell type three separate subsets of nuclei, performing DEA on each, and retaining only DEGs appearing in all 3 subsets. Second, we applied DESeq2 DEA model fitting, to exclude genes with low expression dispersion estimates and low counts^42,43^ (Methods).

Using the above DEA pipeline on DA neurons, and stratifying for HIV status (to assess the impact of SUD), we called 51 genes downregulated in SUD+HIV-vs. SUD-HIV-donors (false discovery rate [FDR] p<0.05; log2 fold change [l2fc] > variable values, dependent upon FDR, sample size, and dataset information content^43^; Data S1). These DEGs enriched gene ontology (GO) terms^44^ linked to neuronal function, including axon myelination, presynaptic organization, cell adhesion, and Na^+^ transport (Fig. 2A; Data S2). Notably, SUD+HIV+u donors, while exhibiting significant sharing of these downregulated DEGs with SUD+HIV-donors (by permutation test, p<10^−5^) also showed 82 uniquely upregulated DEGs, which enriched most prominently GO terms for axon growth and presynaptic transmission (Fig. 2B, J, K; Data S2). This observation suggests enhanced sensitivity to SUD in the context of HIV comorbidity, even in the face of viral suppression (for HIV+u donors). In contrast, SUD+HIV+d had 0 DEGs vs. SUD-HIV+d (Figs. 2C, P; Data S1), suggesting that for these two viremic donor groups, transcriptomic alterations of SN DA neurons were primarily driven by factors related to active HIV replication (viremia).

Indeed, this is what we observed when the DEA of DA neurons was stratified for SUD status (to assess the impacts of HIV), with SUD+HIV+d donors having 40 up-/ 38 down-regulated DEGs vs. SUD+HIV-(Fig. 3B). Furthermore, while downregulated DEGs were highly shared between SUD+HIV+d and SUD+HIV+u donors (35 shared, p<10^−5^; Figs. 3A, B), many viremia-specific (HIV+d) effects were identified (56 downregulated and 5 upregulated DEGs that enriched different GO terms; Fig. 3C, Data S2). Importantly, the presence of SUD increased virus-associated transcriptomic dysregulation: DA neurons from SUD-HIV+d and SUD-HIV+u donors consistently showed a much smaller number of DEGs than their SUD+ counterparts (by approximately 1/3; Figs. 3D, E) and in the absence of SUD, HIV+u and HIV+d transcriptomes were indistinguishable (n DEGs = 0, Fig. 3F).

Despite the above SUD-attributable differences, a striking commonality in all HIV+ groupings also emerged from our analyses: DA neurons from all HIV+ groups, regardless of SUD status or VL, showed significant downregulation in *SLC6A3* dopamine and *SLC18A2* monoamine transporter genes (Figs. 3A, B, D, E; Data S1, S2), together with a more variable dysregulation of genes related to neuronal immune responses, presynaptic vesicle release, synaptic signaling, and cell-cell adhesion. From all the above observations, we draw two conclusions. First, the neurogenomic response to HIV infection, both for donors with viral suppression and without, alters fundamental signaling mechanisms in SN DA neurons including dopamine reuptake. Second, SUD comorbidity enhances abnormality in HIV+ donors, driving a stepwise progression in DA neuron dysregulation, in which abnormalities are more severe in SUD+ compared to SUD-donors and enhance the effects of systemic viral replication (Figs. 3C, F).

Next, we ran our DEA pipeline on SN GABA neurons, which produced few DEGs across all comparisons (there were median [interquartile range or IQR] 0 [0, 0.75] GABA DEGs for each of our 6 DEAs, as opposed to 24 [0, 40.75] DA DEGs; compare Figs. 2A-C to D-F and Fig. 3 to Fig. 4). Stratification by HIV status produced only one DEG attributable to SUD: downregulation of *RBFOX1* GABAergic transcription factor in SUD+HIV+u vs. SUD-HIV+u donors (associated with decreased inhibition^45,46^; Figs. 2D-F, L, M). However, stratification by SUD revealed severe HIV-associated GABA neuron transcriptomic dysregulation in viremic (HIV+d) donors with SUD, pointing to reduced inhibitory neurotransmission (Figs. 4B, C; Data S2). We counted 115 HIV-associated DEGs (96 down, 17 up) in SUD+HIV+d donors vs. SUD+HIV-, and 322 DEGs (266 down, 57 up) in SUD+HIV+d vs. SUD+HIV+u donors. There was significant overlap (p < 10^−5^) between these two comparisons, including downregulated DEGs involved in GABA production, glutamatergic G-protein-coupled receptor (GPCR) signaling, homophilic cell-cell adhesion, and action-potential-associated membrane depolarization (Data S1, S2). In addition, there was downregulation of several opioid-related transcripts, including of postsynaptic opioid-binding cell adhesion molecule (OBCAM) *OPCML*, which is downregulated during chronic opioid agonist exposure^47–49^; the closely related neurotrimin (*NTM*); ephrin-B1 receptor (*EPHB1*), which colocalizes and interacts closely with ephrin-B2 receptor^50,51^ as a mediator of *OPCML* signaling^49^; and non-coding RNA *AC073225*.*1*, previously reported to show decreased expression in opioid use disorder (OUD)^52^. These alterations were specific to SUD+HIV+d, given that SUD-HIV+d vs. SUD-HIV-, SUD-HIV+u vs. SUD-HIV-, and SUD+HIV+u vs. SUD+HIV-comparisons completely lacked significant differences in gene expression (n DEGs = 0).

**Fig. 4.**
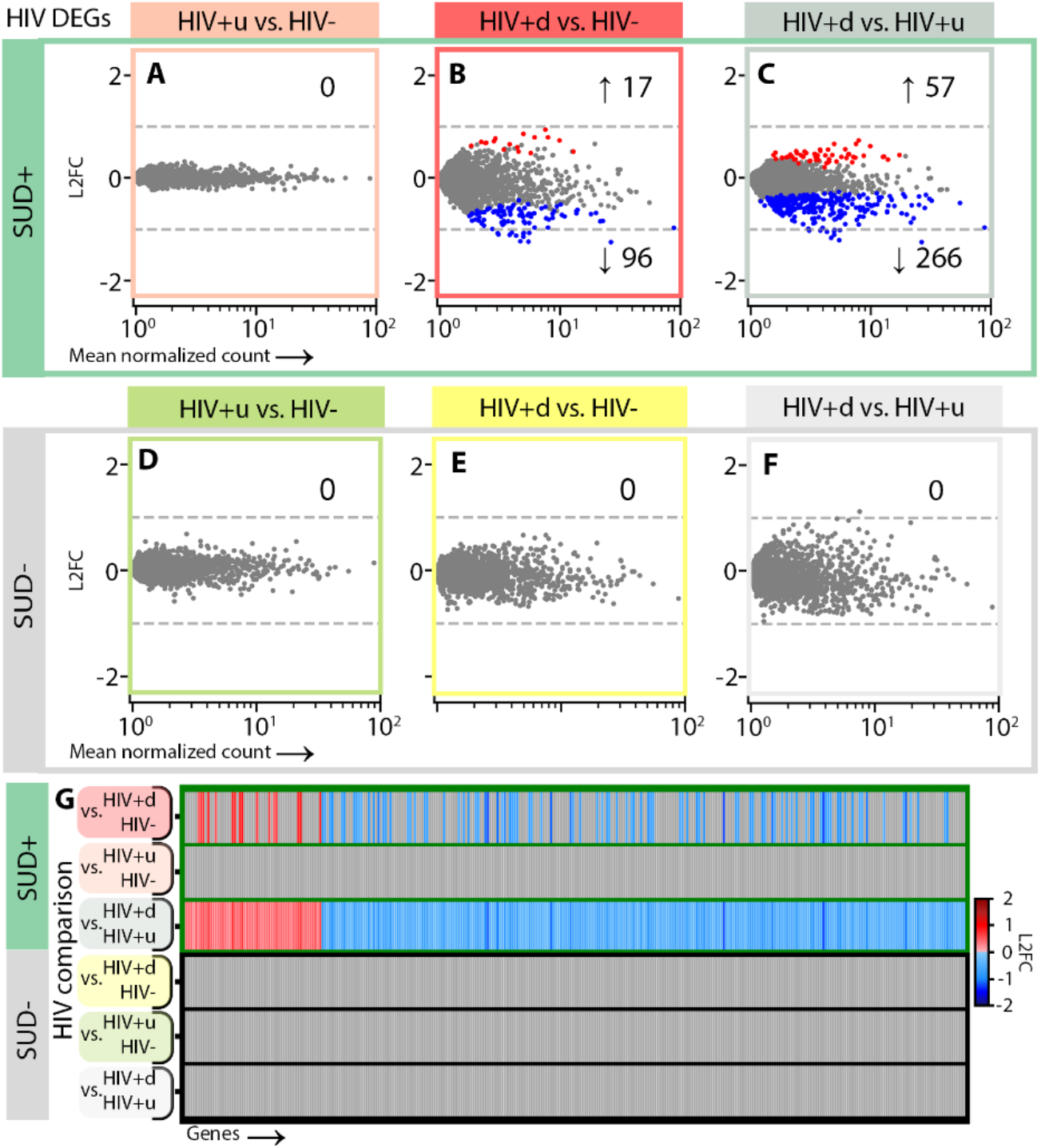
HIV-attributed DEGs for GABA neurons. **(A)-(C)** HIV DEGs (p < 0.05) for GABA neurons in SUD+ donors, for DEAs of **(A)** SUD+HIV+u vs. SUD+HIV-(peach), **(B)** SUD+HIV+d vs. SUD+HIV-(red), and **(C)** SUD+HIV+d vs. SUD+HIV+u (gray). **(D)-(F)** HIV DEGs in SUD-donors for **(D)** SUD-HIV+u vs. SUD-HIV-(green), **(E)** SUD-HIV+d vs. SUD-HIV-(yellow), and **(F)** SUD-HIV+d vs. SUD-HIV+u (gray). **(G)** Gene-by-gene comparisons of differential transcription across HIV comparisons for SUD+ and SUD-donors.

Finally, we applied our stratified DEA pipeline to SN microglia, resulting in hundreds of DEGs attributable to SUD (Figs. 2G-I, N, O, Q). These transcriptional alterations were mostly enriched for various combinations of pro- and anti-inflammatory molecules (Data S1, S2), in broad agreement with recent studies that have reported upregulation of immune signaling genes and transcriptomic signatures indicative of microglial activation and altered glial motility in OUD, chronic opioid exposures, and overdose^34,53–55^. However, in our analysis a substantial fraction of SUD-attributable microglial DEGs were unique to each HIV grouping (Figs. 2N, O), pointing to distinct microglial responses to SUD in the context of varying levels of HIV (HIV-, HIV+u, HIV+d^56^). For example, SUD-associated gene sets supporting reduced protein stabilization and interferon-1 production were seen only in HIV+d, and not in the other HIV groupings (Data S2).

Stratifying by SUD groupings (Fig. 5), we also counted hundreds of microglial DEGs attributable to HIV, including 446 down- and 89 up-regulated DEGs for SUD-subjects with viral suppression (SUD-HIV+u) vs. HIV-controls (SUD-HIV-), >50% of which were replicated in viremic SUD-HIV+d donors (p < 10^−5^). These shared changes primarily involved gene sets for increased inflammatory signaling, decreased phagocytosis and iron ion responses, and increased stress response protein folding/refolding (Data S2). For the SUD-HIV+d group, there were additional changes vs. SUD-HIV+u, including enhancements of immune signaling and telomere maintenance/remyelination processes, as well as antiviral-response-related reductions in glucose transport/GPCR signaling^57,58^. In the context of SUD (Figs. 5A-C), when comparing SUD+HIV+u and SUD+HIV+d donors to SUD+HIV-controls, microglial transcriptional alterations included multiple gene sets related to immune activation, many of which were shared; further, some of these shared effects were opposite to shared HIV effects in the absence of SUD. For example, in the absence of SUD, HIV resulted in downregulated iron ion responses; in the context of SUD, iron ion responses were upregulated (Data S2). Additionally, in the context of SUD, microglia-mediated regulation of synaptic signaling was dysregulated in both HIV+u and HIV+d donors; this effect was absent in the SUD-comparisons.

**Fig. 5.**
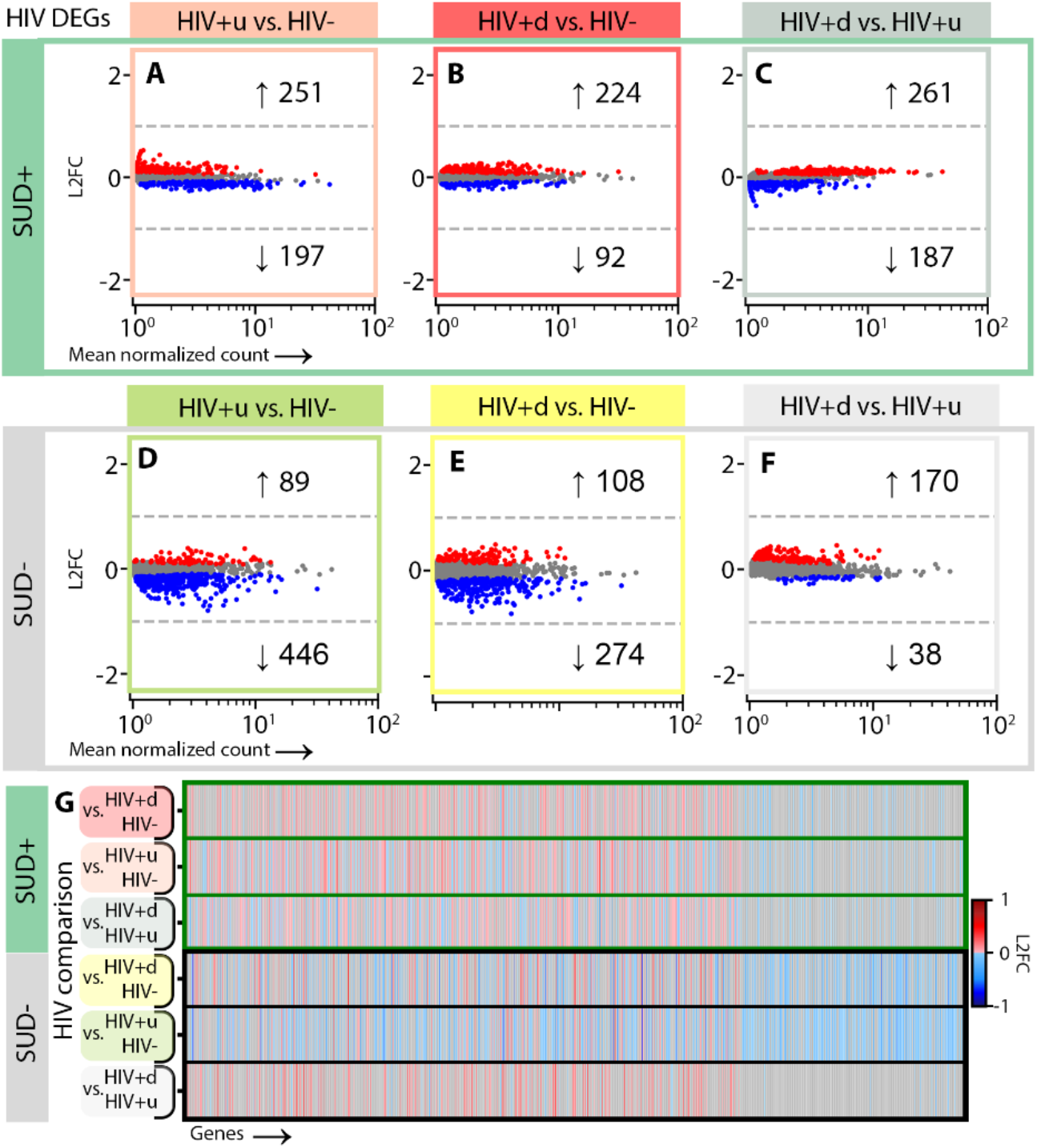
HIV-attributed DEGs for microglia. **(A)-(C)** HIV DEGs (p < 0.05) for microglia in SUD+ donors for DEAs of **(A)** SUD+HIV+u vs. SUD+HIV-(peach), **(B)** SUD+HIV+d vs. SUD+HIV-(red), and **(C)** SUD+HIV+d vs. SUD+HIV+u (gray). **(D)-(F)** HIV DEGs in SUD-donors for DEAs of **(D)** SUD-HIV+u vs. SUD-HIV-(green), **(E)** SUD-HIV+d vs. SUD-HIV-(yellow), and **(F)** SUD-HIV+d vs. SUD-HIV+u (gray). **(G)** Gene-by-gene comparisons of differential transcription across HIV comparisons for SUD+ and SUD-donors.

### Coordinated DEG clusters link microglial and neuronal function, and highlight microglial changes in SUD+HIV+u donors supportive for HIV replication

Next, we wanted to complement our pathway-by-cell type analyses with an alternative approach, by capturing DEGs with statistically linked, or “coordinated”, expression levels across the majority of donors within an SUD/HIV group. To that end, for each DEA we performed, we searched for groups of DEGs that had a high degree of inter-patient covariance (Methods). We selected similarity-based DEG subclusters with average pairwise similarity scores >0.75, our cutoff for identifying gene groups with the most ubiquitous coordinated transcriptional alteration (Fig. 6A), for functional annotation. In the following, we discuss these DEG subclusters, with a focus on microglia-neuron biological functions and interactions.

**Fig. 6.**
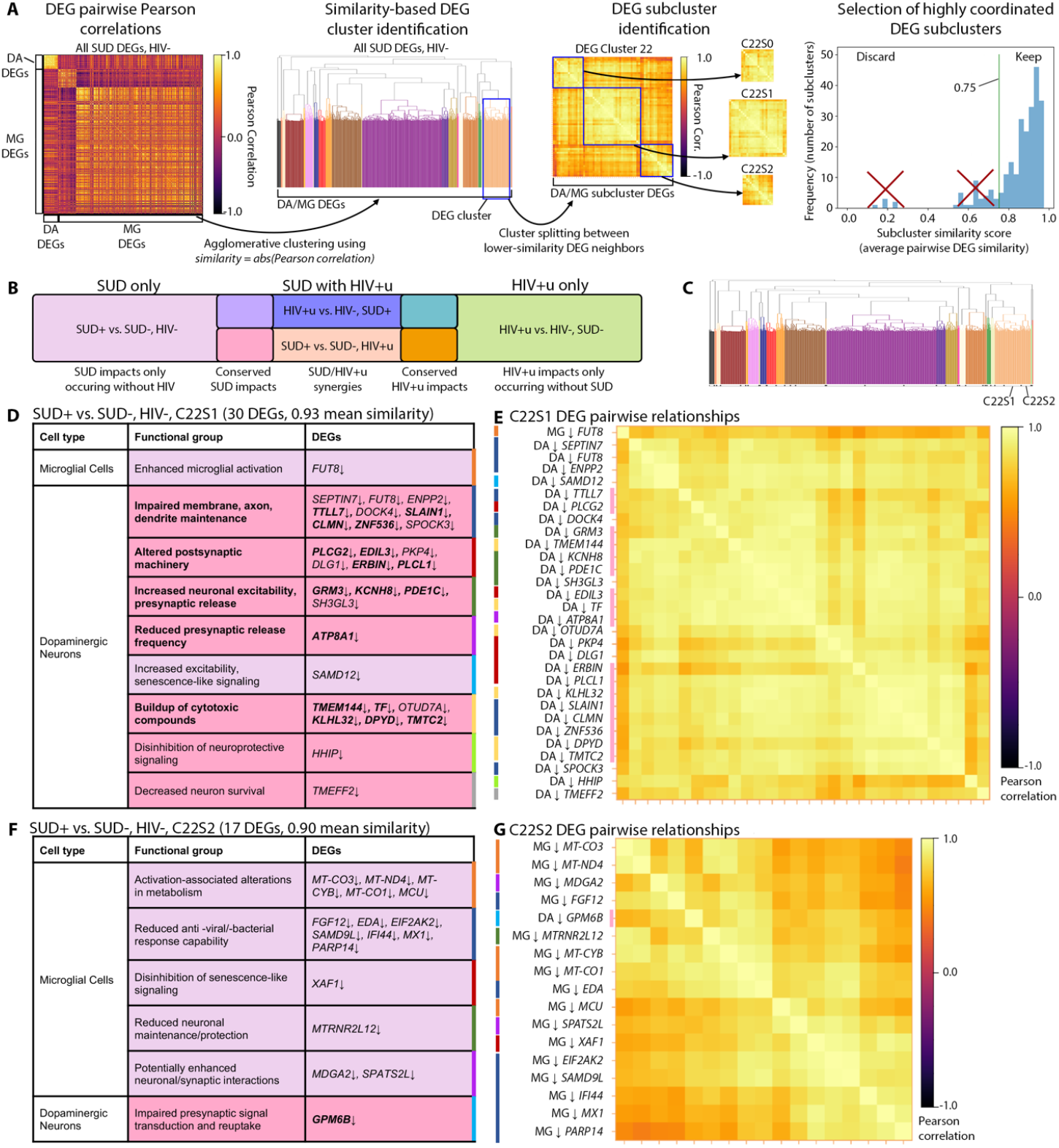
DEG subclustering approach and DA neuron/microglial DEG subclusters showing SUD impacts. **(A)** Schematic of our approach for identifying DEG subclusters, for DA neuron (DA) and microglial (MG) DEGs, for the DEA comparing SUD+HIV-vs. SUD-HIV-(Methods). **(B)** Venn diagram showing interpretations for DEG subcluster functions based on the DEAs they arise in, for the 4 DEAs assessing impacts of SUD (left, **Figs. 2A, G**), HIV+u (right, **Figs. 3D, 5D**), or SUD+HIV+u (middle, **Figs. 2B, H, 3A, 5A**). **(C)** Dendrogram for DEGs from SUD+HIV-vs. SUD-HIV-, showing subclusters for SUD impacts. Labels denote subclusters shown in **(D)-(G). (D)** Subcluster C22S1 DEGs, organized by functional group (indexed by color at right). Fill color follows the legend in **(B)**; bold denotes DEGs recurring as SUD impacts in SUD+HIV+u donors. **(E)** Heatmap showing pairwise DEG correlations for subcluster C22S1, with DEG names colored by functional group to left and pink to right if recurring in SUD+HIV+u. **(F), (G)** Similar to **(D), (E)**, but for subcluster C22S2.

We first examined highly coordinated DEGs for DA neurons and microglia, using the 4 DEAs that showed changes for either SUD, suppressed HIV (HIV+u), or both (Fig. 6B). We identified 117 highly coordinated DEG subclusters (with median [IQR] 10 [6,21] DEGs per subcluster; Table S6, Data S4). Interestingly, 17 of these subclusters contained a mix of DA and microglial DEGs, suggesting DA-microglial interactions. Two of the 17 were specific for the comparison of SUD+HIV-vs. SUD-HIV- (related to impacts of SUD in the absence of HIV; Figs. 6D-G). Both of these subclusters involved glycoproteins: one (C22S1) linked decreased N-glycan synthetic enzyme *FUT8* fucosyltransferase^59^ in both microglia and DA neurons to altered DA neurotransmission (reduced membrane maintenance, reduced pre-/ post-synaptic machinery, slowed presynaptic release, and increased excitability; Figs. 6D, E). Notably, *FUT8* has previously been associated with increased basal microglial reactivity^60^ and impaired neuronal membrane maintenance^61–63^. The other subcluster (C22S2) linked decreased DA neuron expression of *GPM6B* glycoprotein to a microglial switch from oxidative phosphorylation to aerobic glycolysis^64,65^, as well as to deficits in the microglial antiviral response and in transcripts associated with increased microglial-neuroligin-1-mediated excitatory synapse formation (*MDGA2*↓^66,67^) and increased catecholaminergic receptor responses (*SPATS2L*↓^68,69^), among others (Figs. 6F, G).

For dual-diagnosis (SUD+HIV+u) donors, there were 9 mixed DA neuron/microglial subclusters. Three of these closely resembled the subclusters described above for SUD without HIV, specifically for DA DEGs (see “SUD conserved impacts,” denoted by deep pink color, in Figs. 6B, D, F; Data S4). Additionally, and remarkably as these were virally suppressed donors, 5 out of 9 had prominent representation of DEGs linked to HIV infection and replication (Data S4). For example (subcluster C16S2, Figs. 7C, D), DA upregulation of the Na+-Ca2+ exchanger *SLC8A1* was linked to 31 microglial DEGs important for HIV transcription and replication (*HIF1A*↑^70,71^, *HIF1A-AS3*↑^72–74^, *HIVEP2*↑^75–78^, *SYTL3*↑^79,80^, *REL*↑^81–84^, *ELL2*↑^85,86^), HIV host entry and genome stabilization (*TFRC*↑^87^, *SEC14L1*↑^88^, *CD55*↑^89,90^) or viral stabilization (*SIPA1L1*↑ or *E6TP1*↑^91^), and HIV-1 intracellular compartment release (*ATP1B3*↑^92^). We note that Na+Ca2+-exchanger-mediated prolonged neuronal excitation was previously linked to HIV infection in an *in vitro* model^93^. Thus, these subclusters indicated that in virally suppressed PWH, SUD may result in microglial changes permissive for HIV. Other DEG subclusters for SUD+HIV+u donors also linked microglial dysregulation to increased excitability in DA neurons (Figs. 7G, H; Data S4).

**Fig. 7.**
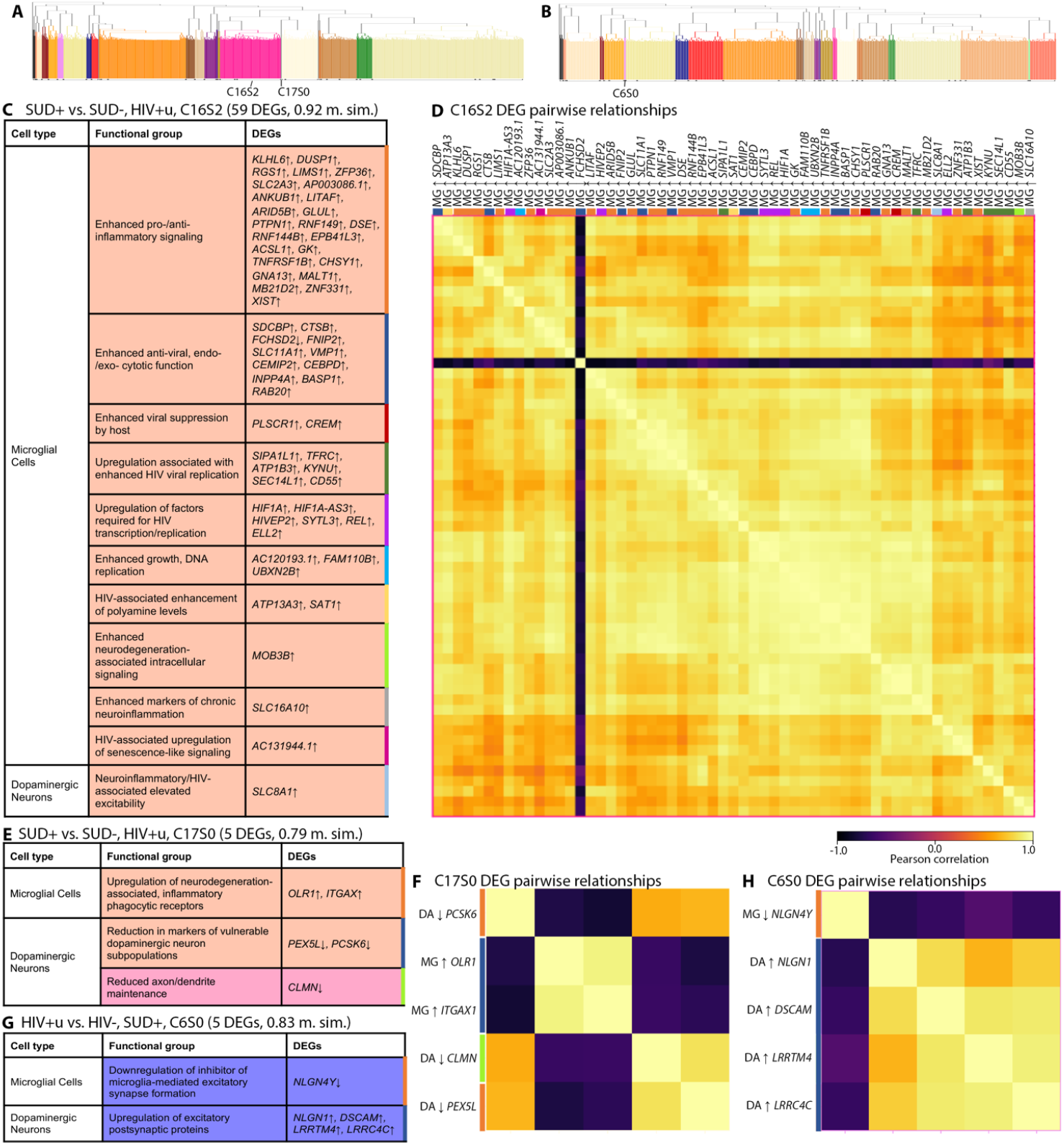
DA neuron/microglial DEG subclusters showing added functional impacts for SUD+HIV+u donors. **(A)** Dendrogram relating DA/MG DEGs for the comparison of SUD+HIV+u vs. SUD-HIV+u (**Figs. 2B, H**), with labels indicating subclusters shown in **(C)-(F). (B)** Dendrogram relating DEGs for SUD+HIV+u vs. SUD+HIV-(**Figs. 3A, 5A**), with labels indicating subcluster in **(G), (H). (C)** Subcluster C16S2 DEGs organized by functional group (indexed by color at right); fill color follows the legend in **Fig. 6B** (pink: SUD impacts; orange: impacts specific to SUD+HIV+u dual diagnosis). “m. sim”: subcluster mean pairwise similarity. **(D)** Heatmap of pairwise DEG correlations for subcluster C16S2, with DEG names colored by functional group. **(E), (F)** Similar to **(C), (D)** but for subcluster C17S0. **(F), (H)** Similar to **(C), (D)** but for subcluster C6S0.

Another specific SUD+HIV+u subcluster (subcluster C17S0, Figs. 7E, F) linked microglial upregulation of receptors for neurodegeneration-associated inflammatory phagocytosis (*OLR1*^94,95^, and *ITGAX*, which targets complement-coated particles^96^) to DA neuronal downregulation of neuronal maintenance genes *CLMN*^97,98^ and *PEX5L*^99^). Taken together, this subcluster suggests that for individuals with undetectable HIV, SUD+/HIV+u synergy still may be associated with damage to vulnerable DA neuron populations.

### In SUD+HIV+d, impaired GABAergic function is coordinated with increased microglial pro-inflammatory signaling

We also identified DEG subclusters for GABA neurons and microglial cells, to explore their relationships in SUD+HIV+d, because in our DEAs this subgroup specifically showed signs of disrupted GABA neuron function (Figs. 4B, C; Data S4). Compared to SUD+HIV+u, SUD+HIV+d individuals had microglial upregulation of DEGs linked to pro-inflammatory signaling (*EPSTI1*^100^, *IFI44L*^101,102^) together with alterations in GABAergic regulators of aging (*RANBP17*↓^103^), cellular maintenance, GABAergic transmission (*SYT9*↓^104^; *DDAH1*↓^105^), and ER-Ca2+-mediated apoptosis (*DPP10*↑, *RYR3*↑^106,107^). Therefore, the impairment of GABAergic regulation in SUD+HIV+d is linked to pro-inflammatory responses in microglia.

## Discussion

We have explored cell-type-specific transcriptional changes in the SN of donors with HIV, opioid/cocaine SUD, or both. Our rationale for HIV groupings (HIV-, HIV+u, HIV+d) was based on an extant literature that has linked plasma HIV to brain microglial activation and other aspects of brain pathology^108^, and on the relevance of plasma HIV load as a standard for guiding patient care. Our rationale for using a polysubstance (opioid/cocaine) approach to SUD groupings was based on the pronounced polysubstance prevalence both in the MHBB cohort and overall in the United States^109^, the importance of these particular drug classes to the HIV epidemic in New York City^27–29^, and a call in existing literature for more research on polysubstance users, which reflects the lifetime exposures of the majority of individuals with chronic SUDs^109–111^.

Our findings, summarized in Fig. 8, demonstrated altered expression of hundreds of pro- and anti-inflammatory transcripts in microglia attributable both to SUD and HIV, with the combined effects of SUD and HIV resulting in a more pronounced dysregulation of the microglial inflammatory transcriptome, particularly in those donors lacking viral suppression (HIV+d). These changes were significantly coupled with abnormalities in neuronal integrity, excitability, and neurotransmission. Perhaps most strikingly, we observed that dysregulated DA neuron transcription in the SN of virally suppressed, opioid/cocaine-SUD+ HIV+u donors was linked to upregulated microglial transcription of genes permissive for HIV infection, replication, and reactivation, along with an altered microglial antiviral response and increased neurodegeneration-associated inflammatory phagocytosis (Data S4). These findings, in a carefully phenotyped, prospectively characterized human population, are striking given the previously reported evidence of a selectively expanded brain viral reservoir in ART-treated, Simian Immunodeficiency Virus (SIV)-infected non-human primates with chronic morphine exposure^112,113^. In this SIV model, reservoir expansion occurred in conjunction with microglial transcriptional changes indicative of immunosuppression and degeneration, relative to SIV+ opioid-negative comparators^112,113^. With the techniques utilized in the present study, we are unable to comment on the brain viral reservoir in our donors, and we identified only 2 HIV transcripts in our entire collection of nuclei (Methods). This is not surprising, given that in a previous study with similar snRNA-seq methodology, HIV transcripts in a subset of microglial nuclei were limited to donor brains displaying the histopathology of active brain viral replication or HIV encephalitis (HIVE)^114^. None of the donors in our present study had histopathologic evidence of HIVE, assessed as part of the standard MHBB neuropathology protocol^115,116^. The lack of viral transcript detection constitutes a limitation of our present work. Furthermore, it contrasts with recent study in which HIV RNA was detected in whole-cell, sorted brain microglia from 2 of 3 virally suppressed HIV+ donors^117,118^. The donors in this prior study were exclusively male and described as having been on ART “until at least 2 wk before death”^117^. The discrepancy between our study and this previous study warrants further investigation, and it could reflect differences in nuclear vs. cytoplasmic substrates for analysis; the time at which ART therapy was discontinued in donors prior to death; or possibly either geographical variabilities in circulating and evolving HIV strains or differing genetic determinants of immune function and viral control in donor populations.

**Fig. 8.**
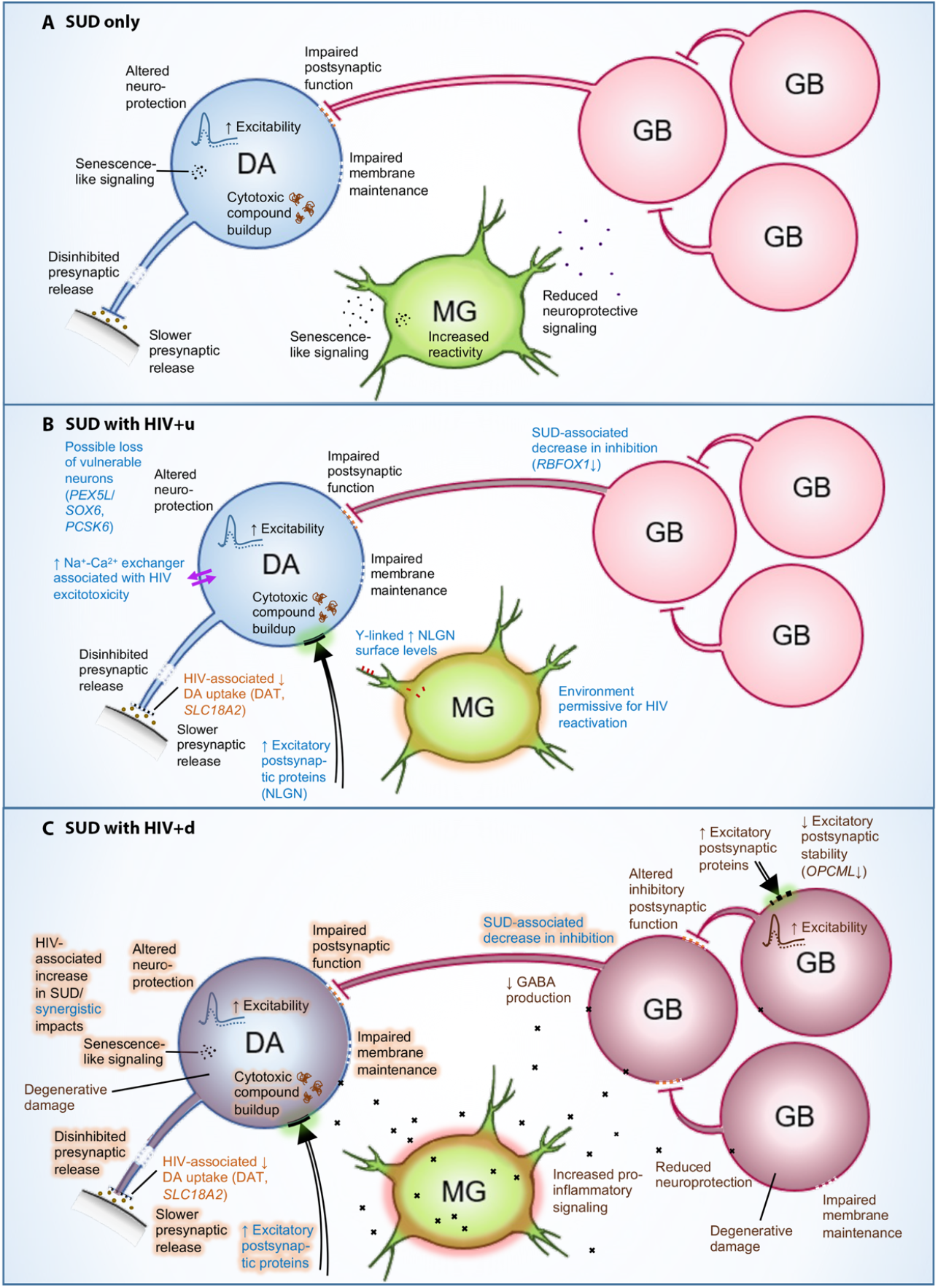
Summary of functional impacts identified for SUD and HIV. Schematics summarizing the biological impacts identified for **(A)** SUD only, **(B)** SUD with HIV+u, and **(C)** SUD with HIV+d, on DA neurons (blue cells; “DA”), GABA neurons (pink cells; “GB”), and microglia (green cells; “MG”). Descriptions indicate biological impacts, with font color showing DEA-based association with HIV and/or SUD (black: SUD impacts; light brown: HIV+u impacts; blue: HIV+u/SUD+ synergistic impacts; dark brown: HIV+d impacts), and with light brown highlights indicating HIV+d-associated enhancement of impacts.

Another important aspect of our work was the demonstration that SUD comorbidity in the context of HIV status exerted impacts on SN neuronal transcriptomes in a stepwise progression. In HIV-donors, we observed only downregulation of DA neuronal transcripts that potentially induce impaired function, senescence-like changes, and increased excitability (Fig. 6, Data S4). In virally suppressed individuals (HIV+u), we observed both a shared (with HIV-) downregulation of DA neuronal genes potentially increasing DA neuron excitability, as well as uniquely upregulated DEGs that suggested enhanced sensitivity to SUD (Fig. 2). Similarly, we observed evidence of enhanced sensitivity to HIV in SUD+HIV+u donors (Fig. 3). With viremia/absence of viral suppression (HIV+d), we further observed changes affecting hundreds of GABAergic transcripts and potentially compromising inhibitory neurotransmission. As microglial transcriptional dysregulation had unique components and also progressed with each HIV grouping, this raises the question of what drives progression, and whether it is unidirectional (driven by one cell type) or bidirectional (driven reciprocally). Recent work in mixed cell cultures (containing microglia with either DA or GABA neurons) has shown that healthy neurons may drive HIV into latency in microglia; and conversely, neuronal damage, including substance-related damage, fosters microglial HIV reactivation and viral expression, thereby further compromising neuronal health due to enhanced microglia-mediated neurotoxicity^119^. Our findings in postmortem human brain raise the possibility that similar bidirectional phenomena may be relevant in human, but future studies will be needed to further examine the mechanisms driving neuronal and glial dysfunction, expanding on our observations of transcriptomic signatures of coordinated detrimental interactions between HIV and SUD (for a more complete listing of significant coordinated abnormalities, see Data S3, S4).

Finally, while we have focused on effects of HIV and SUD comorbidity in altering SN neuronal and microglial transcriptomes, it is noteworthy that in all our HIV+ donor groups, regardless of viral load or SUD status, DA neurons showed a consistent deficit in the expression of the dopamine and monoamine reuptake transporters *SLC6A3* (encoding the protein “DAT”) and *SLC18A2*, respectively, two key regulators of DA turnover at synapses. Combination cocaine- and-raclopride positron emission tomography (PET) studies of viremic PWH, both with and without SUD, have demonstrated similar abnormalities at the protein level *in vivo*, showing decreased DAT signal in the dopamine-terminal-rich basal ganglia^120,121^. The prevailing hypothesis for this phenomenon was that HIV viral proteins, including its transactivator of transcription or Tat and others, directly bind to monoamine transporters, thereby inhibiting monoamine (including dopamine) reuptake and damaging DA neurons^122,123^. However, downregulation of DAT and monoamine reuptake transporter transcripts occurred in our study even in the virally suppressed HIV+ SUD-donor group; and furthermore, evidence of viral Tat transcripts in our brains was lacking. The DA-neuron-specific RNA results presented here point to a transcriptional mechanism resulting in decreased monoamine transporter expression that, while linked to some unified aspect of HIV infection, does not require evidence of ongoing viral replication or transcription.

In summary, the strengths of our study include: a large number of carefully phenotyped brain donors with large amounts of prospectively collected data on SUD and immunovirologic status; novel analytic pipelines that are tolerant to abnormality yet rigidly subjected to quality assurance/QC procedures that exceed other published norms; analytic groups parsed by clinically relevant factors in HIV and SUD; and in-depth annotation of DEG groups that show coordinated expression in most members of their associated donor group (Data S3, S4). Limitations include a lack of viral transcript detection and the lack of mechanistic exploration of the multiple transcriptomic abnormalities described in these donor groupings. Future studies are indicated to further understand the significance of these genomic adaptations for future development of therapies for PWH and SUD.

## Methods

### Brain donor tissue and clinical data collection

Brain donors were autopsied and their donations curated by the MHBB between the years 1999 and 2021, using study protocols and consent documents approved by the Icahn School of Medicine at Mount Sinai (ISMMS) Institutional Review Board (IRB; IRB Approval Number STUDY-11-00388). Briefly, at the time of autopsy, the entire brain was removed, photographed, and bisected, and one hemisphere was placed in phosphate-buffered formalin for subsequent processing, gross neuropathologic review, sectioning, and histological analysis^115^. The other hemisphere was immediately sectioned, each section photographed, and then was snap-frozen between aluminum plates chilled to a temperature of -85° Celsius (C). Frozen brain sections were stored at -85°C in a monitored facility. For this study, samples of ventral midbrain containing recognizable pigmentation indicative of SN were dissected, using a rotary saw on a cold table (chilled to -20°C). As there was some rostro-caudal variation in the anatomical level at which each donor’s SN was dissected, we accounted for the relative location as rostral midbrain (SN at the transverse level of the red nucleus) or caudal midbrain (at the level of the cerebellar peduncular decussation) and included this as a covariate in our differential expression analysis (see below).

Prospectively collected clinical information was available for 67 donors (62 HIV+, 5 HIV-) as a consequence of their participation in the MHBB longitudinal, observational study. For 23 other HIV-donors, information was extracted by medical record review at the time of demise. Substance use characterization (DSM IV dependency diagnoses) for those followed in the MHBB study was obtained with either the Psychiatric Research Interview for Substance and Mental Disorders (PRISM) versions 1.9b and SL^124^ or the World Health Organization Composite International Diagnostic Interview (CIDI) version 2.1^125^, and drug utilization was monitored by urine toxicologic analysis for opiates, cocaine, and other psychoactive substances at each study visit, as described previously^126^. At the time of urine toxicology, the donor’s reported medications were examined to determine whether psychoactive substances were prescribed secondary to medical illness. For donors not enrolled in the prospective MHBB study, DSM IV diagnoses and problematic utilization were obtained from medical records, inclusive of urine toxicologic analysis when available. Immunovirologic data (CD4 T-cell counts and plasma HIV VL) were similarly obtained either as a prospective research study laboratory or retrospectively extracted from medical records; all were performed in the CLIA-certified laboratories of the Mount Sinai Hospital. In this study, for HIV+ individuals, we used a VL of 50 copies/mL as the threshold for HIV detectability, to be consistent with the quantitative lower limit of the earliest assays utilized in the MHBB cohort.

### snRNA-seq

We performed sample processing consisting of nuclei purification, RNA extraction, and subsequent generation of snRNA-seq libraries, following almost exactly a previously described method^34^ for sequencing of pooled tissue samples. Briefly, we homogenized tissue aliquots containing pooled ventral midbrain from SN using a douncer in a mixture of lysis buffer and RNase inhibitor. In each library, we pooled tissue from 1 to 4 unique donors to increase sequencing throughput; we also added a small volume of mouse tissue to most libraries (28/33) as a negative control to create a threshold for spurious HIV reads (for future work). On pooled tissue, we used a lysis and sucrose buffer gradient to isolate nuclei during ultracentrifugation. We resuspended nuclei in a BSA solution containing RNase inhibitor, stained them with DAPI nucleophilic dye, and used fluorescence-activated nuclei sorting (FANS) to collect 300,000 DAPI+ nuclei for input into the 10x Chromium NEXT GEM Single 3’ v3.1 (Dual Index) Protocol according to manufacturer instructions. We quantified libraries using qPCR prior to sequencing on the Illumina NovaSeq platform. Sequencing was performed by the New York Genome Center. As described below, each library was separated by donor after sequencing and prior to analysis, using genotype-based demultiplexing, such that we performed our analyses on a per-donor basis (see below).

We generated the first round of snRNA-seq libraries by pooling tissue from 3 to 4 patients per library, then performed QC and generated additional snRNA-seq libraries for donors with <1,000 QC-passing nuclei, in each pooling tissue from only 1 to 2 patients to increase the per-patient nuclear yield. We added libraries for low-yield donors until the tissue sample was exhausted or the total number of quality-passing nuclei exceeded 1,000 for 90% of donors, resulting in a total of 51 libraries across the 90 study donors. We determined the threshold of 1,000 QC-passing nuclei per donor using a power analysis (see below).

### Imputed donor SNP arrays for demultiplexing

We used donor SNP arrays as a reference to demultiplex pooled snRNA-seq data, generating them from donor cerebellum tissue using the Illumina Infinium Global Screening Array platform (GSA-24v3.0). We prepared DNA for each donor at a final concentration of 20 ng/μL. Genotyping was performed at the Center for Applied Genomics Core at Children’s Hospital of Pennsylvania (CHOP). We performed QC on SNP arrays using GenomeStudio v2.0^127,128^, retaining patient genotypes with call rate of >95% of SNPs, and SNPs with minor allele frequency (MAF) >0.2, call rate >99%, and clustering (“GenTrain”) score >0.7. We performed a second round of SNP genotyping for three donors with SNP call rates <95%, resulting in two panels with 517,091 SNPs (614,679 variants total) and 416,166 SNPs (603,685 variants total) across chromosomal designations 1-22, X, Y, XY, PAR, and MT. To create a demultiplexing reference, we retained SNPs from chromosomes 1-22 for each panel, and we imputed larger arrays for these to improve demultiplexing accuracy^30^ using the Trans-Omics for Precision Medicine (TOPMed) Imputation Server^129–131^ (TOPMed r2 reference, EAGLE v2.4 phasing^132,133^, no *r*^*2*^ filter, data-TOPMed reference panel allele frequency comparisons, pre-run QC). After pre-run QC, 482,995 and 212,783 SNPs from the two panels were used for phasing and imputation; after imputation the panels had 27,749,087 and 11,426,093 SNPs. We combined these imputed genotypes for donors into a single reference and ran demuxlet on each group of sequenced libraries we produced as described above (4 total). For each group, we filtered the demultiplexing reference using bcftools^134^ to retain only biallelic SNPs that had a depth of coverage across donors >9, as well as estimated imputation accuracy *r*^*2*^ >0.5 and MAF >0.2. Our resulting reference panels included 568,817; 480,835; 676,322; and 291,170 SNPs.

### Alignment and cleaning of snRNA-seq data

We pre-processed sequenced data using multiple steps, first performing alignment, demultiplexing, and expression noise/doublet removal on each pooled library separately, then performing heuristic-based filtering on the data from all libraries combined, to generate per-donor data cleaned of ambient RNA, doublets, and other low-quality information.

#### 1. Read alignment

We first aligned reads per library to a composite genome using Cellranger version 6.1.2 (using *count* with option *--include-introns*)^135,136^. We built the composite genome reference using CellRanger’s *mkref*, to be a combination of the GRCh38 human genome, the mm10 mouse genome, and a split version of the HIV-1 reference genome (previously shown to reduce ambiguities in alignment of HIV-associated reads, increasing detection^137^). Notably, we only detected two HIV reads in our data, likely related to both the presence of highly mutated HIV dissimilar to the reference sequence and lower viral levels in our chronically infected, non-encephalitic HIV+ donors (Discussion).

#### 2. Demultiplexing/multidonor doublet removal (demuxlet) + expression noise removal (CellBender)

Second, we performed two processes in parallel, demultiplexing and ambient RNA/noise removal, using as input for each the human-genome-aligned reads for each library. We demultiplexed data with demuxlet^30^ and popscle (https://github.com/statgen/popscle), using our donor SNP reference panel described above, doublet mixing fraction *α* ranging from 0 to 0.5, and 0.1 doublet prior, which produced for each nuclear barcode a label of “singlet”, “doublet”, or “ambiguous” and its maximum-likelihood donor ID(s). We confirmed that for each library the donor IDs returned matched those expected.

We performed ambient RNA/noise removal using CellBender remove-background^138,139^ with the *remove-background-v3-alpha* workflow on the Terra.bio platform^140,141^. CellBender uses unsupervised learning to model and remove expression count noise, due to ambient RNA molecules and random barcode swapping. For most libraries we ran CellBender using false positive rate 0.01, learning rate 0.00005, posterior batch size 5, 150 epochs, and 2 retries. For a few libraries, we adjusted these: CellBender failed to converge during learning for library SNr11 with our original parameters but achieved reasonable loss indicative of successful learning with learning rate 0.000025 over 200 epochs. Another library (SNr27) showed signs of overfitting that we corrected by using learning rate 0.00005 over 60 epochs. For each library, the output from CellBender was a cleaned expression matrix in anndata^142^ format.

We combined demuxlet and CellBender results by filtering each library’s CellBender-cleaned expression data to retain only demuxlet-assigned, nuclear singlet barcodes, then adding the corresponding donor IDs to the expression data as barcode annotations.

#### 3. Multi-cell-type doublet removal (Scrublet)

Third, we used Scrublet^143^ to remove remaining doublets (barcodes containing reads from multiple cell types from one patient; we expected that demuxlet removed most multi-patient doublets), processing each patient’s nuclei within each library separately (134 patient groups across 51 libraries). Scrublet removed 4,372 doublets in total, or a median (IQR) of 33 (8, 37) (1.45% [0.82%, 2.22%]) of doublet barcodes per group. After this step, we presumed that most barcodes corresponded to single nuclei and subsequently called them “nuclei”.

#### 4. Heuristic filtering

Fourth, we combined expression data across all libraries and performed heuristic filtering to remove potentially low-quality or dying nuclei. We filtered based on fraction of mitochondrial genes/nucleus, number of genes/nucleus, and number of reads (denoted by unique molecular identifiers or “UMIs”)/nucleus. For each heuristic, we produced the distribution of values across all dataset nuclei, then identified the thresholds of specific distribution tails and filtered out nuclei with more extreme values. We excluded nuclei with the highest mitochondrial fractions from our dataset, since fractions >5% or 10% are typically thought to indicate potentially dying cells^31^. We removed nuclei in the upper tail of our dataset, amounting to 10% of our nuclei with mitochondrial fractions >1.28% (Fig. S1A). This measure is notably lower than typical thresholds (5% to 10%), such that we may have conservatively over-removed nuclei in this step. We also removed nuclei with excessively high numbers of genes/nucleus (which may indicate e.g. doublets), excluding the upper tail or the ∼5% of our nuclei with >4,396 genes/nucleus (Fig. S1B). Importantly, because previous studies have shown that human SN neurons and some immune cells exhibit lower gene counts in snRNA-seq datasets^31,32^, we did not filter out nuclei with lower genes/nucleus at this step. Instead, we relied on cell-type-specific filtering performed during differential expression analysis to exclude low-count data (see below). Following a similar rationale, we excluded nuclei in the upper tail of our dataset in terms of number of UMIs/nuclei, amounting to ∼5% of our nuclei with >17,153 UMIs (Fig. S1C). Notably, we identified heuristic thresholds each time a new group of sequenced data was added to our dataset (during data generation; see above) and identified roughly the same thresholds as those above during each intermediate stage.

### snRNA-seq data dimensionality reduction, batch correction, and nucleus clustering

To produce a finalized dataset for analysis, we performed three additional steps. First, we further reduced expression count noise using two forms of linear dimensionality reduction: highly variable gene (HVG) selection and principal component analysis (PCA) dimensionality reduction (Figs. S1E, F). Second, we performed batch correction, to facilitate cross-library dataset integration (Data S5). Third, we performed nucleus clustering to use as input for cell typing. Below, we explain several choices that we made in these steps to accommodate expected transcriptional abnormalities for donors. Many of these and other downstream analyses were performed using scanpy^144^ and other packages as noted.

When selecting HVGs (scanpy.pp.highly_variable_genes() with “seurat_v3” option, which ranks genes by cross-nucleus variance^33^), we found that the top 2,000 HVGs excluded many marker genes we used for cell typing, and included genes, like inflammatory markers, that might relate more to cell state. To retain more cell type marker genes but still reduce noise, we ordered all 36,601 genes in our data by increasing HVG rank (decreasing expression variance) and inspected the simultaneous decrease in minimum expression variance and increase in number of cell type marker genes as we considered more HVGs (Fig. S1E). We selected a number of HVGs to retain that was past the elbow of the “number of cell type marker genes” curve, because beyond this point, including additional HVGs would introduce cell type marker genes at slower rates at the cost of adding progressively lower-variance data (a diminishing return). We kept 20,000 HVGs, which preserved 67% of our database’s cell type marker genes (Data S6). Notably, working with this number of HVGs posed a substantial memory burden, since several scanpy functions by default convert expression counts to high-precision data types (64-bit). To alleviate this burden, we created modified versions of several scanpy functions for downstream analysis that maintained data at a user-specified precision; we then found the smallest data precision that would accommodate our expression data (32-bit) and used it as input to those functions.

We performed PCA on HVG-selected expression data (scanpy.tl.pca()) and retained the first 25 components (Fig. S1F) for batch correction and also UMAP (Uniform Manifold Approximation and Projection)^145,146^ generation, clustering, and other downstream analysis. We preserved PCA-reduced, batch-corrected expression data for downstream analysis by projecting it back to the barcodes-by-genes basis from the truncated PCA basis (details below). To facilitate this last step, we stored the means and standard deviations used to scale data prior to PCA as well as PCA loadings using a data-type-modified version of scanpy.pp.scale(), with zero centering and no expression count capping (to recover the original counts data in the limit of no PCA truncation or batch correction).

We ran batch correction using Harmony^147^ (snRNA-seq library ID batch variable, 20 maximum iterations); Harmony converged in 3 iterations. We confirmed that batch correction results were reasonable by comparing pre-vs. post-Harmonization 2D UMAP localizations for nuclei that were from the same patient but processed in different libraries (Data S5).

Batch-corrected data in the PCA basis was used to make a UMAP-based neighborhood connectivity graph using scanpy.pp.neighbors() (20 neighbors) for local manifold approximation; we then generated 2D UMAPs using scanpy.tl.umap() and initial Leiden clusters for cell typing using scanpy.tl.leiden() (resolution parameter 0.5).

We projected the batch-corrected, scaled expression data back to the gene basis from the PCA basis using the 25 stored PCA loadings, then made the resulting scaled expression data count-like, ensuring non-negative counts, then rescaled it using stored means and standard deviations (Fig. S4). Batch-corrected distributions sometimes had non-negligibly negative values (indicative of barcodes with minimal, effectively zero, HVG counts relative to others). Most of these values were between -1 and 0 (Fig. S4B), and any adjusted count values of <-1 occupied <1% of any given HVG’s corrected distribution (Fig. S4C). We thus set these negative values to zero; our approach for doing so was to zero the lowest 1% of each HVG’s count distribution. This approach had no effect on existing non-negative count values because a substantial percentage of each HVG’s distribution was zero counts. We then shifted each distribution so its minimum matched its uncorrected counterpart and would produce non-negative values upon rescaling. Finally, we rescaled the values, set any remaining negative values to zero, and converted to integer-type.

### Likelihood-based cell typing

Because we observed indications of abnormal transcription, we wanted to avoid assumptions about cell types that were present in our data or the most appropriate marker genes to use for cell typing. We thus developed an approach for cell typing based on the principle employed in ScType, a computational cell typing tool^148^. ScType estimates cell types using a large, multi-tissue, multi-organism database of cell-type-specific “positive” and “negative” marker genes that are expressed in or suppressed in a cell type, respectively. For a group of unidentified nuclei or cells and specified tissue(s), ScType computes a “cell type score” for each nucleus and possible cell type, which is a weighted sum of the nucleus’s standardized expression levels across the possible cell type’s positive and negative marker genes. Weights each reflect a “marker gene specificity score”, reducing the impact of less-specific marker genes, and have negative sign for negative marker genes to penalize the score when these are expressed; ScType assigns the cell type with the maximum score to each nucleus.

Our approach, like ScType’s, compares gene expression levels against a cell type marker gene database, but in other respects it differs considerably. We perform cell typing on nuclear clusters rather than individual nuclei, with the reasoning that clusters, by definition, group nuclei with similar transcriptional profiles, and that considering them may highlight genes with characteristic expression levels that may be more helpful for cell typing. We identify such “distinguishing HVGs” for each cluster as the HVGs that are differentially overexpressed vs. all other clusters combined, using scanpy.tl.rank_genes_groups() with *k*-fold, leave-one-out cross-fold validation (*k*=10) on scaled (scanpy.pp.normalize_total()), log1p-transformed (scanpy.pp.log1p()) data (Wilcoxon-rank-sum comparison, Benjamini-Hochberg p-value correction). We considered genes that were overexpressed across all *k*=10 calculations with log2 fold change (l2fc) >1.0 and adjusted p <0.01 to be distinguishing HVGs, and we used these (as opposed to all HVGs) for comparison of the nuclear cluster against our cell type marker gene database.

The cell type database we use in this study is a modified subset of the ScType database^148^ that contains marker genes for relevant human tissues (brain, immune tissue, and smooth muscle tissue, which has cell types resembling those in the blood-brain-barrier). Some ScType cell types appeared in multiple tissues, so we removed any redundancies in these, either merging types with identical marker gene lists or assigning tissue-specific names to cell types with differing lists (e.g. renaming “Endothelial Cell” to “Brain Endothelial Cells”). We then added marker genes from recent postmortem human SN snRNA-seq studies^34,149,150^ to the database. We also retained cell types not expected for SN (e.g. glutamatergic neurons, fibroblasts, cancer cells) to detect potential sample issues and to better characterize unexpected phenotypes. Finally, although we pulled positive and negative marker genes from ScType and have included an option to use them in our code, we only used positive marker genes in this study, to avoid assuming that negative markers would be absent for cells in non-normative states. The final database that we used in our analysis here has 61 cell types, with median (IQR) 14 (6, 27) positive marker genes per cell type, and 515 positive marker genes total (Data S6).

In our approach, we use likelihoods to assess whether a cluster is of a particular cell type. Explicitly, for a cluster *n* of nuclei, and cell type *c*, we define the likelihood *L* as the joint probability that each of *c*’s markers, 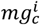 (where *i* is an identifying index running from 1 to the total number of marker genes *M*) is a true, defining marker gene for *n* (meaning that it is in the set of *n*’s distinguishing HVGs, which we denote as *mg*_*n*_):

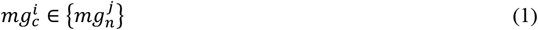

(where *j* is another identifying index). We assume that the probability of 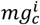 being a marker for *n* is a monotonically increasing function of its overexpression level in *n*, which we denote as 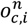. By this, we mean that if 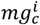 was not overexpressed in *n*, its probability of being a marker for *n* is ∼0, and if it is overexpressed, its probability of being a marker increases with increasing 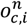, saturating at ∼1 above some overexpression level:

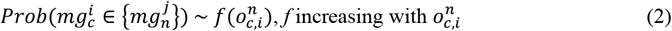

We make an approximation of linear dependence on 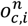,

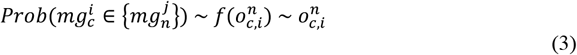

and we make the simplifying assumption that the probability functions for 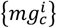 can reasonably be treated as independent. The likelihood then takes the form

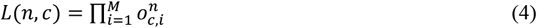

In our calculations we use the log-likelihood *LL*:

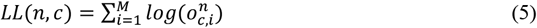

If negative marker genes were to be used (they are not in this study), the log likelihood becomes

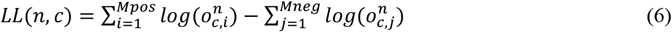

We do not apply cell-type-specificity or other weighting to marker gene likelihood terms, to avoid inadvertently making biased assumptions. For example, by not applying weights, we avoid inappropriately upweighting marker genes that appear for fewer database cell types as a result of missing data rather than biological specificity. We also avoid discounting less-specific marker genes that may nevertheless be informative for cell typing.

We construct the log-likelihood in Eq. (5) by setting 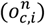, the log overexpression for marker gene *i* of cell type *c*, to either the mean (across *k* calculations) l2fc expression of that gene in cluster *n* if the gene was a distinguishing HVG (and thus significantly overexpressed), or 0 if not (overexpression level 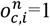). Using this measure, we observe substantial drop-off in log-likelihood as the overlap between a cluster’s distinguishing HVG’s and a cell type’s marker genes decreases (Fig. S5), leaving only a few putative cell types. We used the highest-likelihood cell types and the contributing database marker genes for each to assign a cell type to each nuclear cluster, and we used this approach in multiple rounds to arrive at final cell types. First, we ran cell typing on our original Leiden clusters, then combined contiguous clusters with the same cell type, then performed subclustering on any of the resulting clusters that had visible substructure (8 total), again using a Leiden algorithm but with resolution 0.1 on each cluster in isolation. Subclustering had the effect of splitting nuclei groups with different cell types that were originally merged, including microglia and macrophages, and GABAergic and dopaminergic neurons. Finally, we performed cell typing again on these re-clustered nuclei, generating the cell types in Fig. 1. We found our cell types to be in agreement with previously published human SN snRNA-seq data^34^ (Fig. S3).

### Differential expression analysis (DEA)

We performed Wald-formulation DEAs on each cell type separately with *k*-fold cross-validation (*k*=3 set by power analysis; below), using DESeq2 ^43^ ported into Python with the package diffexpr (https://github.com/wckdouglas/diffexpr/tree/master). For SUD+ vs. SUD-comparisons, the generalized linear model (GLM) we used as DESeq2 input was

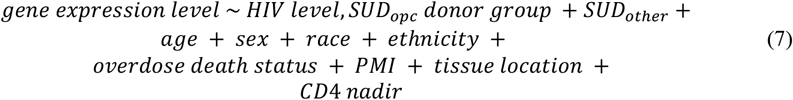

For HIV comparisons (HIV+u vs. HIV-; HIV+d vs. HIV-; HIV+d vs. HIV+u), our GLM was

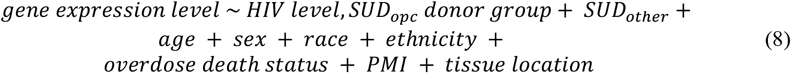

Covariates are defined as follows:

*HIV level*: Categorical. Final HIV viral load in copies RNA/mL. (Options are “HIV-”, “HIV+u” for <50 copies/mL, “HIV+d” for >50 copies/mL)

*SUD*_*opc*_: Categorical. Presence of opioid and/or cocaine SUD diagnosis. (“yes”, “no”) *SUD*_*other*_: Categorical. Presence of SUD diagnosis for substance other than opioids/cocaine. (“yes”, “no”)

*Age*: Numeric. Donor age in years.

*Sex*: Categorical. Donor sex assigned at birth. (“female”, “male”)

*Race*: Categorical. Donor race described in clinical records. (“Black or African American”, “White”, “Asian”, “Black and White”, “Black and Indian”, “Unknown or Not Reported”) *Ethnicity*: Categorical. Donor ethnicity described in clinical records. (“Hispanic or Latino”, “Not Hispanic or Latino”)

*Overdose Death Status*: Categorical. Presence of overdose as cause of death. (“yes”, “no”)

*Postmortem Interval (PMI)*: Numeric. Donor post-mortem interval in hours.

*Tissue Location*: Categorical. Approximate anatomical location of SN sample. (“coronal” for rostral midbrain, “midbrain” for caudal midbrain)

*CD4 Nadir*: Categorical. Lowest donor CD4 T-cell count. (“<200 cells/µL”, “[200,500] cells/µL”, “>500 cells/µL”). HIV-donors are assigned “>500 cells/µL”.

Prior to running DESeq2, numerical covariates were standardized, and categorical covariates that covaried perfectly were identified, and all but one were removed, to avoid ambiguity in fitting. None of the DEAs in this study required us to drop covariates.

DESeq2 implements multiple controls to remove false positives^43^, and we used *k*-fold cross-validation as an additional layer of false positive control for each DEA. In *k*-fold cross-validation, random subsetting of data is expected to randomly alter low gene dispersions. Low dispersions can exist for genes with prominent within-donor correlations (e.g. if there are many similar nuclei per donor relative to the rest of the DEA data) and/or low counts, and they can inflate the apparent effect size of expression changes, leading to false positives. The randomly altered dispersions across *k*-fold cross-validation computations alters the degree to which such genes are false positives. We filtered to include only DEGs that show up in all *k* random subsets (below), which helps to remove these impacts. DESeq2 can also be run on single-nucleus data rather than pseudobulked data (for which nuclei are aggregated per cell type and donor); the single-nucleus approach can help to highlight DEGs that may be pertinent to subsets of a cell type if the nuclei exhibit biological variability^42^, and for this reason we ran DESeq2 on our single-nucleus data without pseudobulking. DESeq2 authors have made recommendations for run parameters that increase the number of “true positive” DEGs returned when running on single-nucleus data by handling zero inflation characteristic of single-cell approaches^42^; we have confirmed that those recommendations reproduce our results and add additional DEGs. Importantly, our procedure is more restrictive in terms of DEGs returned but still benefits from false positive controls and does not produce DEGs not seen when run with DESeq2-author-recommended parameters. One comparison is shown below for DEA results using our parameters (Wald test, useT=False, default sizeFactors, and pseudocount addition) vs. zero-inflation recommendations (likelihood ratio test, useT=True, minmu=1e-6, sizeFactors from scran::computeSumFactors).

| Cell type | DEA | N up DEGs reported in our paper | N up DEGs common to both approaches | N down DEGs reported in our paper | N down DEGs common to both approaches |
| --- | --- | --- | --- | --- | --- |
| DA | SUD+HIV- vs. SUD-HIV- | 0 | 0 | 51 | 49 |
| DA | SUD+HIV+u vs. SUD-HIV+u | 82 | 76 | 41 | 41 |
| DA | SUD+HIV+d vs. SUD-HIV+d | 0 | 0 | 0 | 0 |

To perform DEA, we formed *k* data subsets from a cell type’s nuclei, then fit the DESeq2 model to each subset (using the GLM in Eq. 7 when assessing SUD impacts or Eq. 8 when assessing HIV impacts; py_DESeq2() followed by run_deseq()). We then pulled the results for each donor group comparison of interest (e.g. SUD+HIV-vs. SUD-HIV-donor groups) using dds.deseq_result(). With this approach, DEA model fitting for each *k*th subset was based on nuclei from ∼90 donors across all donor groups, such that fits of gene dispersion benefitted from a large sample size and higher nuclear variability. We identified differentially expressed genes (DEGs) as those with adjusted p-values <0.05 across all *k* calculations. For significant DEGs, we used the cross-fold means of l2fc and normalized counts in Figs. 2-5 and for gene set enrichment analysis.

We included an extra term in our SUD GLM (Eq. 7), for CD4 T cell nadir, that was not in our HIV GLM (Eq. 8). This is because, although we split HIV groups by the clinically relevant measure of final VL, we noted that other lifetime characteristics of HIV infection could impact transcription, most notably long-term damage incurred during periods of low T-cell counts. We thus added CD4 nadir to Eq. 7 to account for longer-term HIV impacts, and to aid detection of SUD impacts. We found that CD4 nadir tracked closely with final HIV VL for our HIV+ donors, accounting for a small degree of HIV-related variation. For HIV comparisons, we excluded CD4 nadir because it closely recapitulated HIV VL variation.

### Gene set enrichment analysis (GSEA)

We performed GSEA with Enrichr^44^ using GSEApy, a Python wrapper for R-based GSEA tools^151^. For each DEA, we ran Enrichr separately for up- and down-regulated DEGs (gseapy.enrichr()), using our 20,000 HVGs as background and human annotated gene set reference “GO_Biological_Process_2023”^152,153^. We saved all gene sets returned regardless of significance but considered gene sets with adjusted p-value <0.05 to be significantly endorsed (Data S2). We also tested the tool GSEA^154,155^ in GSEApy, and found that it identified similar but generally fewer significant gene sets for DEGs (Table S7, Data S9).

### Identification of highly coordinated DEG subclusters

To find DEG subclusters for a particular DEA and set of cell types, we first pulled the normalized count data (scanpy.pp.normalize_total(); as opposed to l2fc or any other DEA result), for all nuclei of each cell type of interest, across all that DEA’s case group donors. We then computed for each cell type, each DEG’s mean normalized expression count (across all that cell type’s nuclei) per case donor. This step produced a matrix of cell-type-specific DEGs per row, and the mean normalized expression of that DEG for each case donor per column. To distinguish DEGs that arose for multiple cell types, and to keep track of the direction of dysregulation, we named DEGs using the convention c*ell-type_DEG-direction_gene-name*. For all DEGs across all cell types, we then computed cross-donor Pearson correlations in mean expression for DEG pairs using scipy.stats.pearsonr(), using the built-in beta-assumption hypothesis test to determine significance (two-tailed alternative) with significance level 0.05. Across all our computations (Table S8) there were median (IQR) 30% (28%, 34%) DEG pairs with significant correlations.

We computed the similarity for each DEG pair to be the correlation magnitude if it was significant or 0 if not significant; for DEGs *i,j*

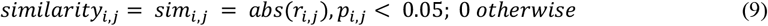

To use hierarchical agglomerative clustering to group DEGs, we transformed similarities to distances using

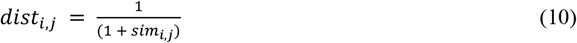

adding 1 to the similarity to avoid division by zero. We used Eq. 10 instead of *dist*_*i,j*_ = 1 − *sim*_*i,j*_ because its nonlinearity required DEG clusters to have more pronounced similarity to be detected, by inflating distances between higher-similarity DEG pairs, and thus by suppressing the dynamic range of distances used to parse clusters. We performed clustering using sklearn.cluster.AgglomerativeClustering() (average linkage distance) with “distance_threshold”=0 and “n_clusters”=None to compute the entire linkage tree and return algorithm-generated DEG clusters. The resulting DEG clusters often contained noticeable subclusters, with inter-subcluster distances substantially larger than intra-group distances. We thus split them at points where the distance between adjacent DEGs was ≥3 standard deviations above the cluster’s mean adjacent DEG distance. To keep track of original cluster identity, we named DEG subclusters using the convention C*[cluster-index]*S*[subcluster-index]* (e.g. “C17S2” for cluster 17, subcluster 2). We characterized subcluster coordination strength by computing the average similarity across all DEG pairs in a subcluster. Some subclusters had exceptionally low average similarities (∼0.2 vs. ∼0.8; Fig. 6A, right), indicating weaker overall relationships with other DEGs. To focus on annotating the most highly coordinated DEG groups, we removed these low-similarity clusters by excluding anything in the lower tail of the distribution of average similarities across all DEG subclusters (<0.75; Fig. 6A right).

### DEG subcluster annotations

We produced a skeleton annotation .csv file for each DEG subcluster using “GO_Biological_Process_2023” annotations and the acyclic graph of their relationships (“go-basic.obo”). For each DEG, we found all related GO terms and filtered them to include only the most subordinate annotation represented on each graph branch (most specific process). We split remaining terms into those explicitly mentioning positive regulation of a process, explicit negative regulation, explicit response, and others, as an initial basis for annotation (Data S3). We used DEG naming convention *cell-type_DEG-direction_gene-name* to facilitate interpretation.

For focused DEG subcluster annotations, two experts (MJ, AW) independently searched different gene databases for each DEG (MJ, GeneCards; AW, National Center for Biotechnology Information gene database) and pulled the functional summary. They then performed a literature search for the DEG using terms related to HIV, SUD, brain location (“substantia nigra”, “ventral midbrain”), and cell type (e.g. “gene *x* in neurons”, “gene *x* in dopaminergic neurons”, “gene *x* in microglia”). Relevant references were collected and used to annotate potentially relevant functions for each gene; these results were then used to generate a consensus across annotators that was used in each DEG subcluster (Data S4).

### Statistical analysis

#### snRNA-seq sample size selection

We used a power analysis simulation to estimate the number of nuclei per donor needed to power detection of cell-type-specific expression differences. We simulated a case where expression data was compared across two groups for one gene in one cell type, and there was a small but true difference in the two distributions. We defined each group’s underlying expression distribution for the gene to have a lognormal distribution, i.e.

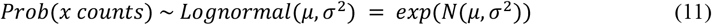

We let both distributions have the same variance of 1 and means differing by some l2fc value. The difference between means *μ*_1_ and *μ*_2_ was related to the l2fc by

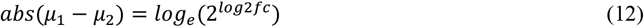

For each l2fc we tested (0.5, 1.0, 1.5), we defined the two underlying distribution means using Eq. 12, then over 10,000 iterations sampled distributions of size *s* (nuclei) from each and performed a Wilcoxon rank sum test of distinguishability. We computed the power as the fraction of iterations in which the true difference was detected (p <0.05). We repeated this simulation starting with *s*=10 and incrementing *s* by 2 until we reached a power ≥80%. We achieved sufficient detection power with *s*=158 for l2fc 0.5; *s*=42 for l2fc 1.0; and *s*=20 for l2fc 1.5. Based on this simplified situation, we set a threshold of ∼100 nuclei per cell type, aiming for detection of expression differences at l2fc 1.0. Prior to analysis, we estimated there would be ∼10 expected cell types in SN (DA and GABA neurons, microglia, astrocytes, ODCs, OPCs, multiple blood-brain-barrier cells, multiple immune cells^149,156^) and thus aimed to sequence 1,000 nuclei per donor.

### Overexpression level and k selection for cluster “distinguishing HVG” detection

We used a similar power analysis to estimate the l2fc at which distinguishing HVG detection was powered for every cluster of nuclei. We set a significance threshold of p=0.01 to restrict detected distinguishing HVGs to a higher-confidence subset, since these could impact cell typing and thus all downstream analysis. We let distributions represent expression of a gene for one cluster vs. the rest, choosing normal distributions with equal variance and differing means to reflect log1p pre-processing (applied for scanpy.tl.rank_genes_groups()). The means and l2fc were related by

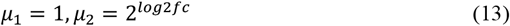

At l2fc=1.0, every cluster exceeded the minimum sample size for 80% detection power. We determined the number of folds *k* for cross-validation by increasing *k* from 2 in increments of 1 until every cluster with one of its *k* folds removed had enough nuclei to also power detection. Our code also had an option for the resulting *k* to be increased by 1, to put each one-fold-removed group of nuclei further over the power detection threshold during cross-fold validation calculations. We enabled this option, finding *k*=9 and using *k*=10.

### Overexpression threshold and k selection for DEA

We used the same power simulation to estimate the values of l2fc and *k* that would power DEAs for each cell type at p=0.05, using *k*=3 and l2fc=1.0.

### Significance testing of cell type proportions across donor groups

We tested cell type proportions were differently represented across donor groups by first using the binomial distribution to model cell type proportions for each donor group. For a donor with *k* nuclei of a specified cell type and *n* nuclei total, the maximum likelihood estimation for the probability of that cell type’s occurrence, 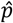, is

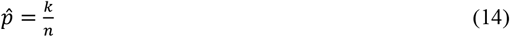

Within each donor group, we computed for each donor the 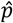 for each cell type, producing a distribution of each cell type’s occurrence probability across group donors. For each cell type, we performed Kolmogorov-Smirnov tests to compare these distributions for each pair of donor groups, using Benjamini-Hochberg multiple comparison correction (12 comparisons, one per cell type, per donor group pair). We considered donor group pairs/cell types with adjusted p <0.05 to have significant differences (Table S5).

### Significance test for DEG overlaps

We performed random sampling tests to determine whether fractions of overlapping DEGs observed for related DEAs (Figs. 2-5) were significantly different from expected outcomes of random sampling. For each DEA pair, we sampled the observed numbers of DEGs per DEA from the 20,000-HVG background, and we recorded the fraction of overlapping DEGs between the two simulated populations. We repeated sampling for 100,000 iterations and computed the upper limit on the p-value to be the fraction of iterations with at least as many overlapping DEGs as observed. In all tests we found 0 such iterations, corresponding to p<10^−5^.

## General

We thank the staff and patients of the Manhattan HIV Brain Bank; this research would not be possible without their valuable contributions. We also thank all members of the Single Cell Opioid Responses in the Context of HIV (SCORCH) consortium for stimulating and helpful discussions, and Roger Pique-Regi for guidance on demuxlet software. This work was supported in part through the computational and data resources provided by the Scientific Computing and Data Center at Icahn School of Medicine at Mount Sinai. The content is solely the responsibility of the authors and does not necessarily represent the official views of the National Institutes of Health.

## Author contributions

Conceptualization: SA, SM

Methodology: AW, MJ, SA, SM

Software: AW

Validation: AW, MJ, AV, TL, SA, SM

Formal analysis: AW

Investigation: AW, MJ, GM, EG, JM, AV, TL, SM

Resources: SA, SM

Data curation: AW, MJ, AV, TL, SM

Writing—original draft: AW

Writing—review & editing: AW, MJ, SA, SM

Visualization: AW

Supervision: AW, MJ, GM, SA, SM

Project administration: SA, SM

Funding acquisition: SA, SM

## Data and materials availability

All data needed to evaluate the conclusions in the paper are present in the paper and/or the Supplementary Materials. Data presented in this study were produced as part of the Single Cell Opioid Response in the Context of HIV consortium (SCORCH: RRID:SCR_022600). De-identified snRNA-seq data, donor SNP arrays, and associated metadata will be deposited at dbGaP (https://www.ncbi.nlm.nih.gov/gap/) and made available prior to publication. The data at dbGaP is controlled-access and requires special handling; please follow dbGaP guidelines and contact dbGaP for support. Mapped, demultiplexed snRNA-seq data will also be made publicly available at the NEMO (Neuroscience Multi-omic) Archive under identifier nemo:dat-zngw2c8 (https://assets.nemoarchive.org/dat-zngw2c8) prior to publication. Code used to perform the analyses in this paper will be made publicly available on Github and Zenodo prior to publication.

## Supplementary Materials for

**This PDF file includes:**

Figs. S1 to S5

Tables S3, S6, S8

**Other Supplementary Materials for this manuscript include the following:**

Tables S1, S2, S4, S5, S7

Data S1 to S9

**Fig. S1.**
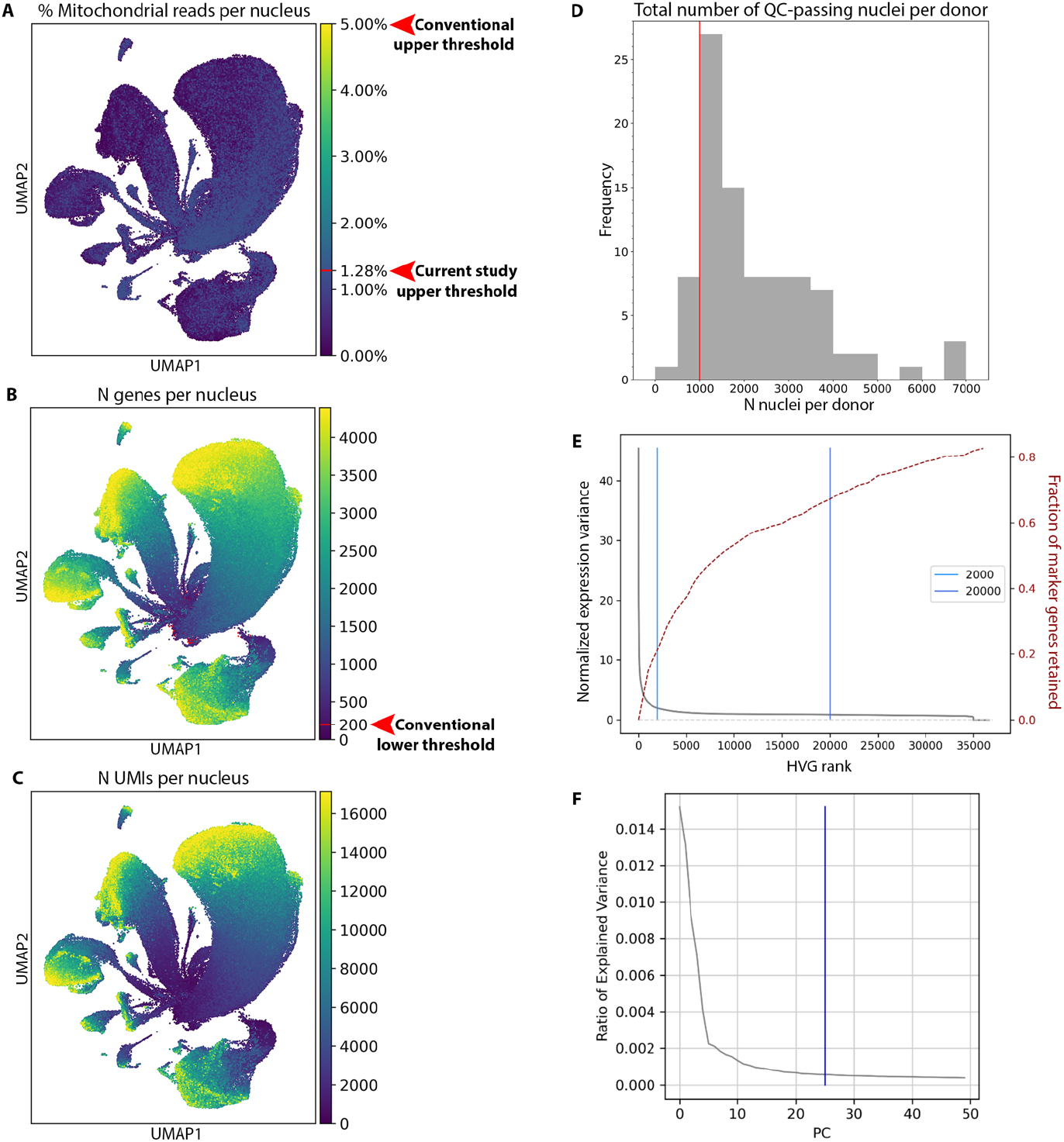
Additional information about snRNA-seq QC and pre-processing. **(A)-(C)** UMAPs showing values of QC heuristics, only for nuclei passing all QC nuclei. **(A)** Percent mitochondrial reads/nucleus. Red line/arrowhead at 1.28% shows the maximum value in our QC-passing nuclei; line at 5% shows a conventional filtering maximum. **(B)** Number of genes/nucleus; red line/arrowhead shows a conventional lower threshold of 200; QC-passing nuclei with <200 genes are colored red (see Fig. S2). **(C)** Number of UMIs/nucleus. **(D)** Distribution of the number of QC-passing nuclei/donor. Red line: the 1,000-nuclei threshold exceeded by 90% of donors. **(E)** Parameters used to determine the number of HVGs retained in analysis. Genes are ranked left to right by decreasing variability; black line (left y-axis) shows each gene’s normalized expression variance; red line (right y-axis) shows the fraction of all genes ≤ that rank in our marker gene database. Blue lines show (right) our cutoff (20,000 HVGs) and (left) a conventional one (2,000 HVGs). **(F)** Explained variances of principle component analysis components (“PCs”); blue line shows the maximum component we retained, the 25^th^ PC.

**Fig. S2.**
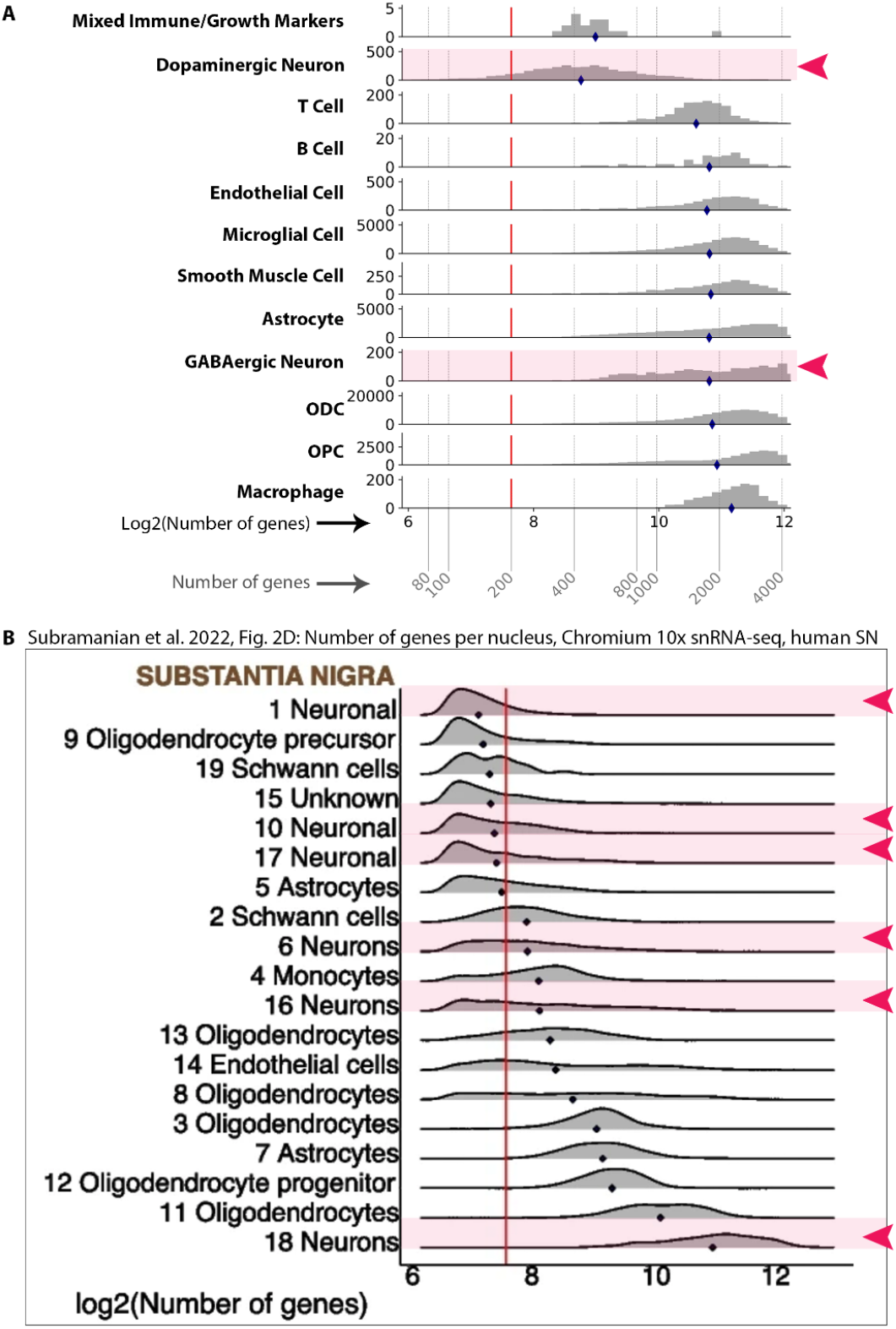
Comparison of N genes/nucleus, by cell type, to published SN snRNA-seq data. **(A)** Histograms showing the log-transformed number of genes/nucleus, by cell type, for our QC-passing nuclei (**Figs. 1B, C**). Gray dashed lines show corresponding raw (non-log-transformed) gene counts. **(B)** Histograms showing log-transformed number of genes/nucleus for each cluster identified in previously published human SN snRNA-seq data, prepared similarly to ours (via Chromium 10x^1^). For **(A)** and **(B)**, neuronal cell types are marked with pink highlights and arrowheads, distribution means are indicated by blue diamonds, and red lines show the conventional lower threshold of 200 genes/nucleus.

**Fig. S3.**
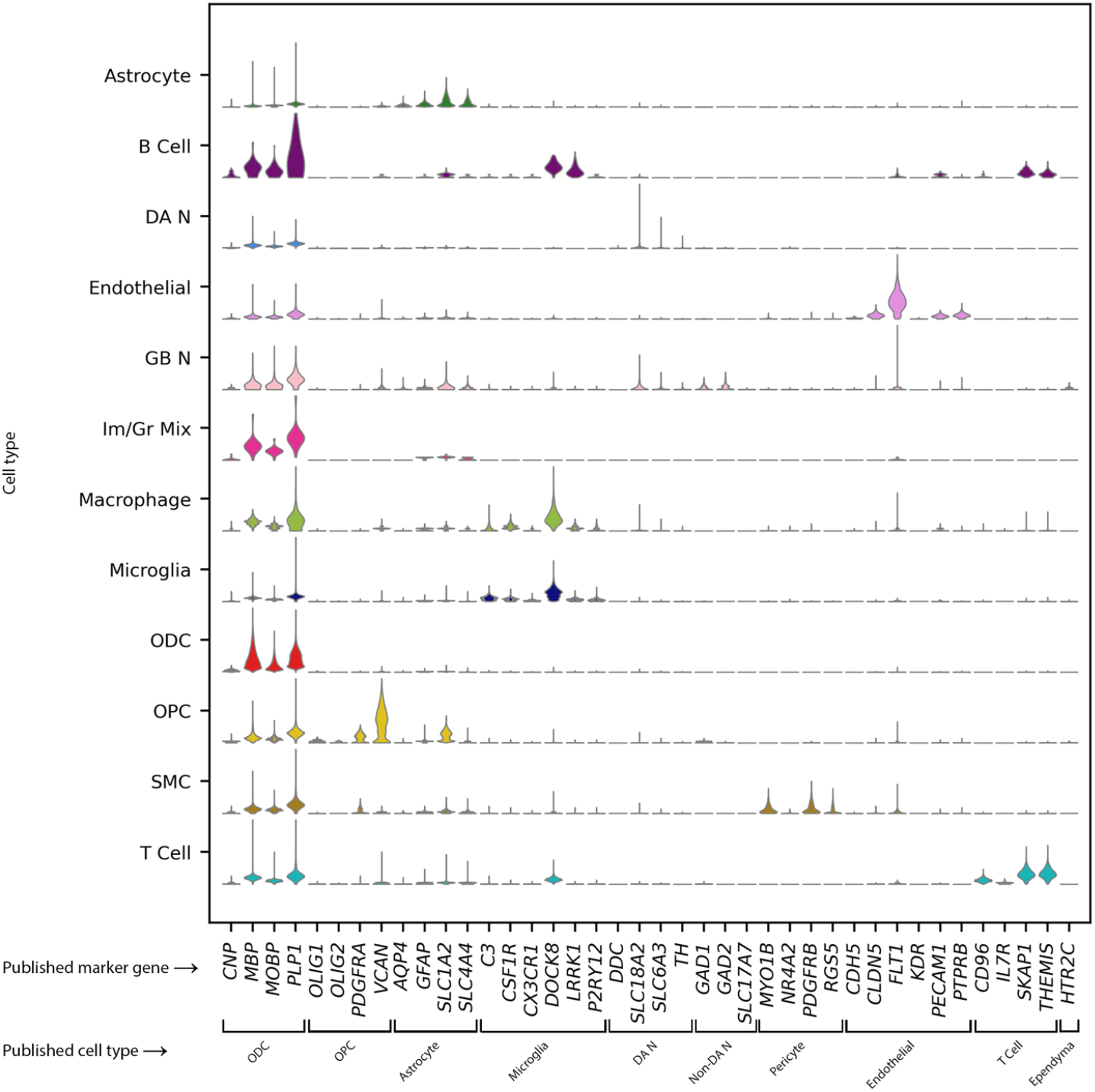
Expression of previously published SN cell type marker genes, by cell type. Expression in each of our cell types (rows) of previously identified (Wei et al. 2023^2^) marker genes for ventral midbrain cell types (columns).

**Fig. S4.**
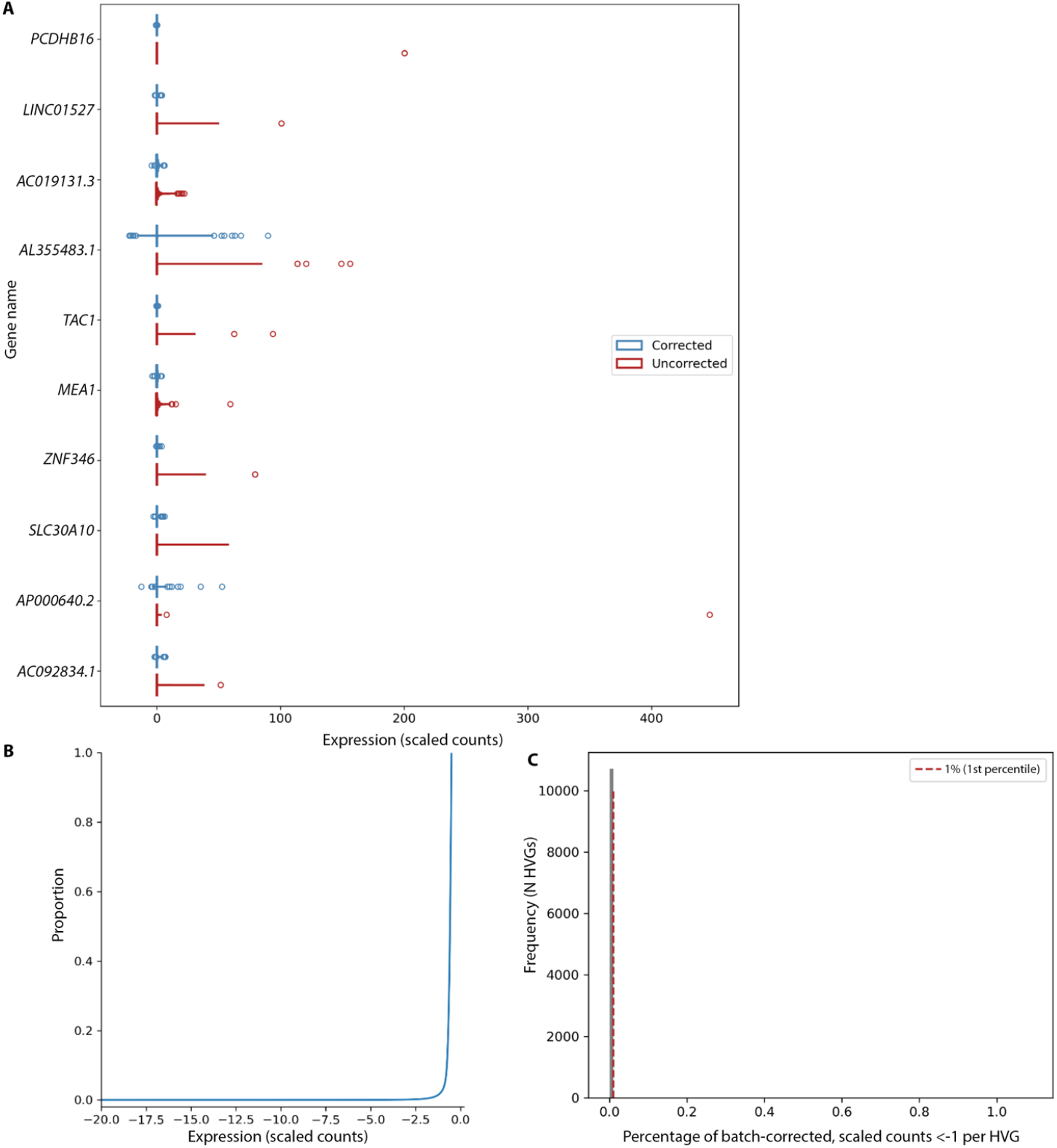
Impacts of batch correction on gene expression distributions for QCed data. **(A)** Gene expression distributions before (red) vs. after (blue) batch correction, for 10 example HVGs selected randomly from our 20,000. **(B)** An empirical cumulative distribution showing the values of all batch-corrected, still-scaled read counts with values ≤ -0.5 (that would round to non-negligible negative counts ≤-1), highlighting a vanishing number of counts shifted to values <-2 by batch correction. **(C)** Distribution showing the percentages of each HVG’s batch-corrected counts with values <-1. Red line denotes 1% (corresponding to the lowest 1^st^ percentile of an HVG’s distribution).

**Fig. S5.**
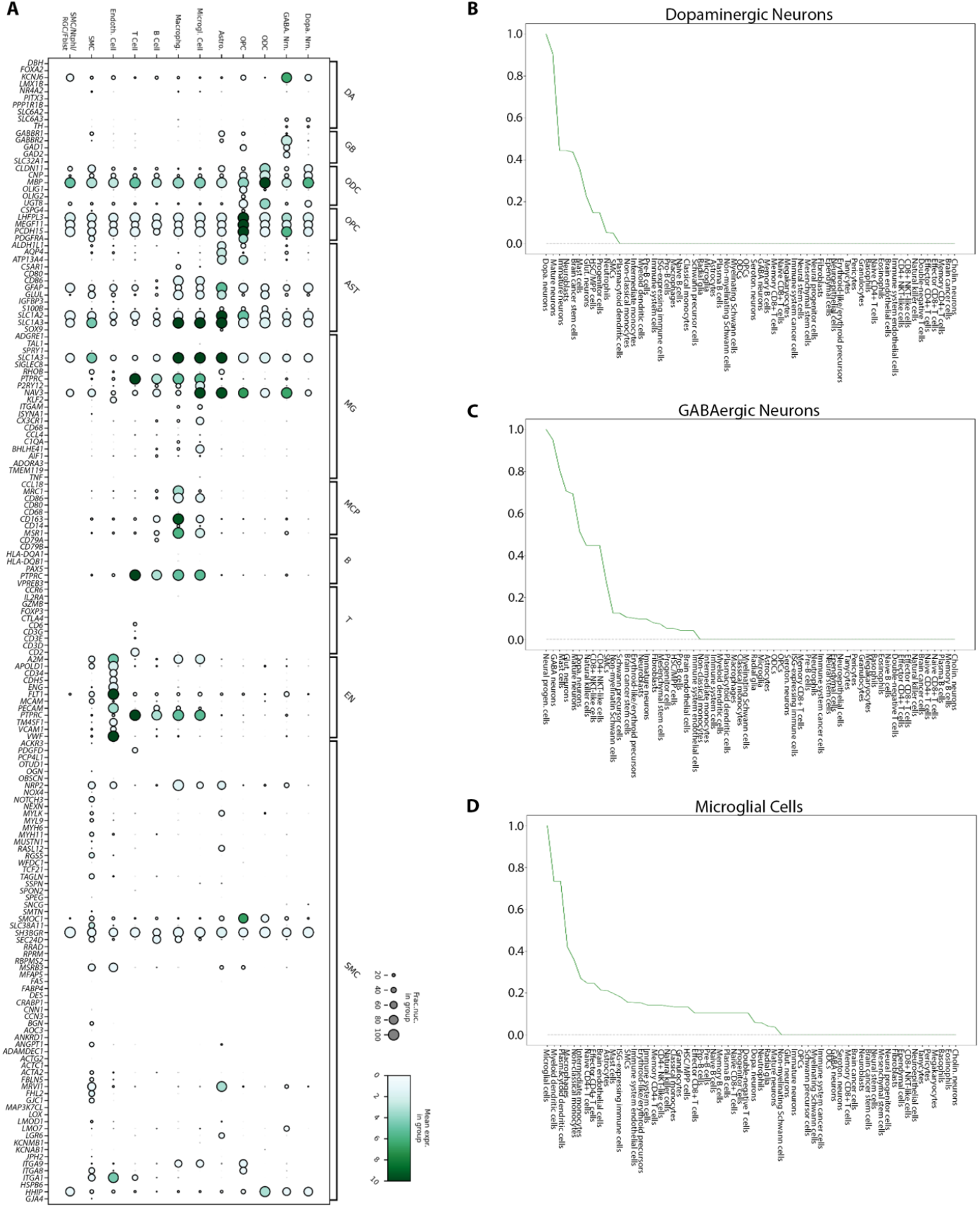
Further information about likelihood-based cell typing of clusters. **(A)** Dot plot showing, for each cell type in our dataset (top labels), expression of various expected SN cell type marker genes (left labels: expected marker genes, right labels: expected SN cell types). **(B)-(D**) Possible database (Data S6) cell types, ordered left to right by decreasing log-likelihood (and normalized to a maximum value of 1), for our **(B)** DA neuron, **(C)** GABA neuron, and **(D)** microglia clusters.

Tables S1, S2, S4, S5, S7

Data S1 to S9

**Table S1. (separate file)**

Donor characteristics.

**Table S2. (separate file)**

Statistical comparisons of donor characteristics across donor groups. Bold text with asterisks indicates p<0.05. Double dashes indicate repeated comparisons or not-relevant self-comparisons.

**Table S3**.

Numbers of donors and nuclei per SUD drug class, by HIV status.

**Table S4. (separate file)**

snRNA-seq nuclear yields by donor and cell type.

**Table S5. (separate file)**

Statistical comparisons of cell type proportions across donor groups. Multiple-comparison-adjusted p-values are shown for Kolmogorov-Smirnov tests; bold and asterisks indicate p<0.05. The cell type proportions per donor group used in these comparisons are shown below this table.

**Table S6**.

DA neuron (DA)/microglia (MG) DEG subcluster compositions, for SUD+/HIV+u DEAs.

**Table S7. (separate file)**

Comparison of gene set enrichment results produced using Enrichr vs. GSEApy.

**Table S8**.

Fractions of significantly correlated DEG pairs in subclustering analyses.

**Table S3.** Numbers of donors and nuclei per SUD drug class, by HIV status.

| **HIV status** | **N donors**<br>**N nuclei**<br>**cocaine SUD** | **N donors**<br>**N nuclei**<br>**opioid SUD** | **N donors**<br>**N nuclei**<br>**cocaine +**<br>**opioid SUDs** | **N donors**<br>**N nuclei**<br>**neither SUD** | **Total N for HIV**<br>**status** |
| --- | --- | --- | --- | --- | --- |
| **HIV-** | 1<br>2,083 | 4<br>5,421 | 10<br>28,610 | 13<br>27,349 | 28<br>63,463 |
| **HIV+u** | 6<br>16,593 | 4<br>10,607 | 10<br>23,281 | 10<br>21,683 | 30<br>72,164 |
| **HIV+d** | 11<br>20,867 | 6<br>19,229 | 8<br>15,333 | 7<br>9,676 | 32<br>65,105 |
| **Total N for**<br>**SUD drug**<br>**class** | 18<br>39,543 | 14<br>35,257 | 28<br>67,224 | 30<br>58,708 | 90<br>200,732 |

**Table S6.** DA neuron (DA)/microglia (MG) DEG subcluster compositions, for SUD+/HIV+u DEAs.

| DEA<br>(effect) | Total N<br>DEG<br>subclusters | Median (IQR)<br>N DEGs per<br>subcluster | Median (IQR)<br>subcluster<br>mean<br>similarity | N mixed<br>DA/MG DEG<br>subclusters | Median (IQR)<br>N DEGs per<br>mixed DA/MG<br>subcluster | Median (IQR)<br>mixed DA/MG<br>subcluster<br>mean<br>similarity |
| --- | --- | --- | --- | --- | --- | --- |
| SUD+HIV- vs.<br>SUD-HIV-<br>(SUD impacts<br>without HIV) | 28 | 13 (7, 23) | 0.88<br>(0.83, 0.94) | 2 | 24 (20, 27) | 0.91<br>(0.91, 0.92) |
| SUD-HIV+u vs.<br>SUD-HIV-<br>(HIV+u impacts<br>without SUD) | 34 | 8 (5, 20) | 0.94<br>(0.89, 0.95) | 6 | 10 (7, 14) | 0.90<br>(0.87, 0.92) |
| SUD+HIV+u vs.<br>SUD-HIV+u<br>(SUD impacts<br>with HIV+u) | 28 | 12 (7, 21) | 0.89<br>(0.85, 0.92) | 6 | 16 (8, 38) | 0.87<br>(0.80, 0.91) |
| SUD+HIV+u vs.<br>SUD+HIV-<br>(HIV+u impacts<br>with SUD) | 27 | 9 (5, 22) | 0.89<br>(0.85, 0.92) | 3 | 9 (7, 23) | 0.90<br>(0.87, 0.91) |
| All | 117 | 10 (6, 21) | 0.91<br>(0.86, 0.95) | 17 | 12 (6, 20) | 0.90<br>(0.86, 0.92) |

**Table S8.** Fractions of significantly correlated DEG pairs in subclustering analyses.

| Cell type mix | N DEAs analyzed | Median (IQR) fraction significantly correlated DEG pairs per DEA, across DEAs |
| --- | --- | --- |
| DA neurons, microglia | 4 | 0.30 (0.28, 0.30) |
| GABA neurons, microglia | 2 | 0.27 (0.27, 0.28) |
| DA neurons, GABA neurons | 2 | 0.46 (0.44, 0.49) |
| All | 8 | 0.30 (0.28, 0.34) |

**Data S1. (separate file)**

DEG lists, organized by cell type (DA neuron, GABA neuron, or microglia) and DEA type (impacts of HIV or SUD).

**Data S2. (separate file)**

Lists of Enrichr gene sets enriched by DEGs per DEA, cell type, and direction of dysregulation (up/down). Significantly enriched gene sets (with adjusted p-value <0.05; column E) are highlighted in green.

**Data S3. (separate file)**

Basic annotations for DEG subclusters, organized by cell type.

**Data S4. (separate file)**

Focused annotations for DEG subclusters, with references.

**Data S5. (separate file)**

UMAPs comparing expression data before (left) vs. after (right) batch correction, showing nuclei from one donor across pooled libraries (“batches”). Each plot title lists the donor’s ID, and each library is shown in a different color.

**Data S6. (separate file)**

Cell type marker gene database used for cell typing.

**Data S7. (separate file)**

Heatmaps of DEG subclusters showing significant DEG pairwise Pearson correlations and related underlying data.

**Data S8. (separate file)**

Heatmaps showing mean normalized counts for each subcluster’s DEGs, per case group donor, with donors labeled by SUD drug class (opioid, cocaine, or opioid+cocaine) and organized by their cross-DEG expression similarities (shown with a dendrogram above each heatmap).

**Data S9. (separate file)**

Zip file containing lists of GSEApy prerank gene sets enriched by each DEA’s DEGs, organized by cell type. Files with names ending in “_all” include all gene sets having any DEGs; files ending with “_sig” include only significantly enriched gene sets.

